# A novel framework for characterizing genomic haplotype diversity in the human immunoglobulin heavy chain locus

**DOI:** 10.1101/2020.04.19.049270

**Authors:** O. L. Rodriguez, W. S. Gibson, T. Parks, M. Emery, J. Powell, M. Strahl, G. Deikus, K. Auckland, E. E. Eichler, W. A. Marasco, R. Sebra, A. J. Sharp, M. L. Smith, A. Bashir, C. T. Watson

## Abstract

An incomplete ascertainment of genetic variation within the highly polymorphic immunoglobulin heavy chain locus (IGH) has hindered our ability to define genetic factors that influence antibody and B cell mediated processes. To date, methods for locus-wide genotyping of all IGH variant types do not exist. Here, we combine targeted long-read sequencing with a novel bioinformatics tool, IGenotyper, to fully characterize genetic variation within IGH in a haplotype-specific manner. We apply this approach to eight human samples, including a haploid cell line and two mother-father-child trios, and demonstrate the ability to generate high-quality assemblies (>98% complete and >99% accurate), genotypes, and gene annotations, including 2 novel structural variants and 16 novel gene alleles. We show that multiplexing allows for scaling of the approach without impacting data quality, and that our genotype call sets are more accurate than short-read (>35% increase in true positives and >97% decrease in false-positives) and array/imputation-based datasets. This framework establishes a foundation for leveraging IG genomic data to study population-level variation in the antibody response.

## Introduction

The immunoglobulin heavy (IGH) and light chain loci comprise the building blocks of expressed antibodies (Abs), which are essential to B cell function, and are critical components of the immune system(*1*). The IGH locus, specifically, consists of >50 variable (IGHV), >20 diversity (IGHD), 6 joining (IGHJ), and 9 constant (IGHC) functional/open reading frame (F/ORF) genes that encode the heavy chains of expressed Abs(*2*). Greater than 250 functional/ORF IGH gene segment alleles are curated in the IMmunoGeneTics Information System (IMGT) database(*2*), and this number continues to increase(*3–10*). The locus is enriched for single nucleotide variants (SNVs) and large structural variants (SVs) involving functional genes(*3, 11–18*), at which allele frequencies are known to vary among human populations(*4, 18, 19*).

Locus complexity has hindered our ability to comprehensively characterize IGH polymorphisms using high-throughput approaches(*20, 21*); only two complete haplotypes in IGH have been fully resolved. As a result, IGH has been largely overlooked by genome-wide studies, leaving our understanding of the contribution of IGH polymorphism to antibody-mediated immunity incomplete(*4, 20, 21*). While early studies uncovered associations to disease susceptibility within IGH, few links have been made by genome-wide association studies (GWAS) and whole genome sequencing (WGS)(*20, 22, 23*). Moreover, little is known about the genetic regulation of the human Ab response, despite evidence that features of the Ab repertoire are heritable(*12, 19, 24–29*)

To define the role of IGH variation in Ab function and disease, all classes of variation must be resolved(*3, 12, 15, 19, 29, 30*). Although approaches have been developed for utilizing genomic or Adaptive Immune Receptor Repertoire sequencing (AIRR-seq) data, variant calling and broad-scale haplotype inference are restricted primarily to coding regions(*6, 7, 16–18, 31*). To fully characterize genetic diversity in the IG loci, specialized genotyping methods capable of capturing locus-wide polymorphism with nucleotide resolution are required. Indeed, such methods have been applied to better resolve complex and hyper-polymorphic immune loci elsewhere in the genome(*32, 33*).

Long-read sequencing technologies have been used to detect chromosomal rearrangements(*34*), novel SVs(*35, 36*), and SVs missed by standard short-read sequencing methods(*37, 38*), including applications in the complex killer immunoglobulin-like receptors (KIR)(*39, 40*) and human leukocyte antigen (HLA)(*34, 41*) loci. Furthermore, the sensitivity of SV detection is improved by resolving variants in a haplotype-specific manner(*38, 42*). When long-read sequencing has been combined with specific target enrichment methods, using either a CRISPR/Cas9 system(*43, 44*) or DNA probes(*45, 46*), it has been shown to yield accurate and contiguous assemblies. Such targeted approaches have enabled higher resolution genotyping of the HLA loci(*47*) and KIR regions(*48, 49*).

Here, we present a novel framework that utilizes IGH-targeted long-read sequencing, paired with a new IG genomics analysis tool, IGenotyper (https://igenotyper.github.io/), to characterize variation in the IGH locus (Fig. 1). We apply this strategy to eight human samples, and leverage orthogonal data and pedigree information for benchmarking and validation. We demonstrate that our approach leads to hiqh-quality assemblies across the IGH locus, allowing for comprehensive genotyping of SNVs, insertions and deletions (indels), SVs, as well as annotation of IG gene segments, alleles, and associated non-coding elements. We show that genotype call sets from our pipeline are more comprehensive than those generated using alternative short-read and array-/imputation-based methods. Finally, we demonstrate that use of long-range phasing/haplotype information improves assembly contiguity, and that sample multiplexing can be employed to scale the approach in a cost-effective manner without impacting data quality.

**Fig 1.**
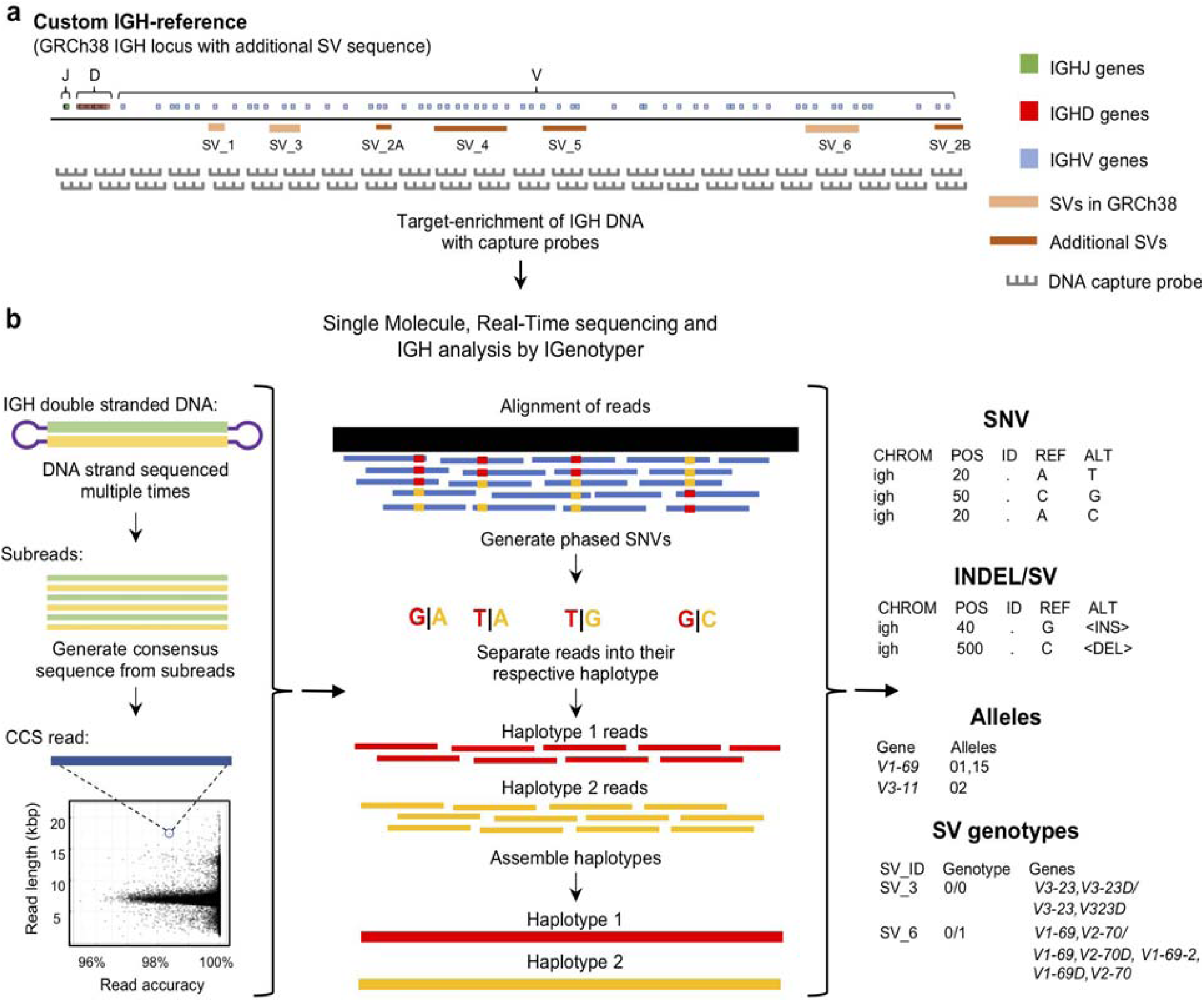
An overview of the custom capture and IGenotyper workflow used to detect IGH variation in a haplotype-specific manner. **a** A schematic of the custom IGH-reference used in this study, which includes the GRCh38 IGH locus with the addition of known SV sequences inserted into their respective positions in the locus. Brown bars indicate the positions of inserted SVs in the IGH-reference; pink bars indicate the positions of additional known SVs present in GRCh38 relative to GRCh37. Positions of IGHJ (green), IGHD (red) and IGHV (blue) genes in the IGH-reference are also indicated. The oligo capture probes used in the panels designed for this study were based on the sequence of this custom IGH-reference. **b** After targeted long-read capture, constructed libraries undergo SMRT sequencing, and the resulting data is processed with IGenotyper. Raw SMRT sequences with at least 2 subreads are converted into higher accuracy CCS reads. Phased SNVs are identified from the CCS reads and used to phase the both the CCS reads and subreads, which are then used to assemble haplotype-specific assemblies of IGH. SNVs, indels, SVs and IG gene/alleles are identified from the assembly and CCS reads. SVs embedded into the IGH-reference are genotyped using the assembly, CCS reads, and SNV calls.

## Results

### Novel tools for comprehensively characterizing IG haplotype diversity

To interrogate locus-wide IGH variants, we implemented a framework that pairs targeted DNA capture with single molecule, real time (SMRT) sequencing (Pacific Biosciences) (Fig. 1a). Roche Nimblegen SeqCap EZ target-enrichment panels (Wilmington, MA) were designed using DNA target sequences from the human IGH locus. Critically, rather than using only a single representative IGH haplotype (e.g., those available as part of either the hg19 or GRCh38 human reference assembly) we based our design on non-redundant sequences from the GRCh38 haplotype(*4*), as well as additional complex SV and insertion haplotypes distinct from GRCh38(*4, 50*) (Fig. 1a; Table S.1). Additional details, including the exact target sequences used and additional specifications of these capture panels are provided in the supplementary material (Supplementary material; Table S.2, 3).

To process and analyze these long-read IGH genomic sequencing data, we developed IGenotyper (Fig. 1b; igenotyper.github.io/) which utilizes and builds on existing tools to generate diploid assemblies across the IGHV, IGHD, and IGHJ regions (see Materials and Methods); for ease, we will henceforth refer to these three regions (excluding IGHC) collectively as IGH. From generated assemblies, IGenotyper additionally produces comprehensive summary reports of SNV, indel, and SV genotype call sets, as well as IG gene/allele annotations. For read mapping, SNV/indel/SV calling, and sequence annotation, the pipeline leverages a custom IGH locus reference that represents known SV sequences in a contiguous, non-redundant fashion (Fig. 1a); this reference harbors the same sequence targets used for the design of target-enrichment panels, and ensures that known IGH SVs in the human population can be interrogated.

### Benchmarking performance using a haploid DNA sample

We first benchmarked our performance using genomic DNA from a haploid hydatidiform mole sample (CHM1), from which IGH had been previously assembled using Bacterial Artificial Chromosome (BAC) clones and Sanger sequencing(*4*). This reference serves as the representation of IGH in GRCh38. Using panel designs mentioned above, we prepared two SMRTbell libraries with an average insert size of 6-7.5 Kb for sequencing on both the RSII and Sequel systems (Table S.4). We observed a mean subread coverage across our custom IGH reference (Fig. 1a) of 557.9x (RSII) and 12,006.4x (Sequel 1), and mean circular consensus sequence (CCS) read coverage of 45.1x (RSII) and 778.2x (Sequel 1). The average Sequel CCS phred quality score was 70 (99.999991% accuracy), with an average read length of 6,457 bp (Fig. S1). Noted differences in target-enrichment panels tested are described in Supplemental Note 1.

To most effectively use these data to assess IGenotyper performance, we combined reads from both libraries for assembly (Table S.4). A total of 970,302 bp (94.8%) of IGH (chr14:105,859,947-106,883,171; GRCh38) was spanned by >1,000x subread coverage, and 1,006,287 bp (98.3%) was spanned by >20x CCS coverage (Fig. 2a). The mean CCS coverage spanning IGHV, IGHD, and IGHJ coding sequences was 160.3x (median=42.5x; Fig. 2b). Compared to GRCh38, IGenotyper assembled 1,009,792 bases (98.7%) of the IGH locus in the CHM1 dataset (Fig. 2c; Table 1). Gap sizes between contigs ranged from 177 to 3,787 bp (median = 456 bp) in length. Only 37 (<0.004% of bases) single nucleotide differences were observed when compared to GRCh38 (base pair concordance >99.99%). In addition, 220 potential indel errors were identified (Table S.5). The majority of these (199/220) were 1-2 bp in length, 61.8% of which (123/199) occurred in homopolymer sequences, consistent with known sources of sequencing error in SMRT sequencing and other NGS technologies (Table S.6). We also observed a 2,226 bp indel consisting of a 59mer tandem repeat motif near the gene *IGHV1-69* (Fig. S2). This tandem repeat was unresolved in GRCh38 (Fig. S2, Supplementary material), which was reconstructed using a Sanger shot-gun assembly approach(*4*); it remains unclear whether this event represents an improvement in the IGenotyper assembly over GRCh38, or is a sequencing/assembly artifact. Nonetheless, the total number of discordant bases associated with indels (2,521 bp) accounts for only <0.28% of the assembly.

**Table 1.** Assembly statistics and evaluation of the accuracy of the haplotype-specific assemblies

| Sample | Contigs (n) | Assembly size (bp) | Assembly validation |  |  |
| --- | --- | --- | --- | --- | --- |
|  |  |  | Concordance with fosmids (SMRT sequencing) | Concordance with BACs or fosmids (Sanger sequencing) | Concordance with Pilon/Illumina |
| CHM1 | 16 | 1,026,385 | NA | 99.996% | NA |
| NA19240 | 38 | 1,829,616 | 99.996% | 99.99% | 99.99% |
| NA12878 | 45 | 1,442,310 | 99.995% | 100.0% | 99.99% |

**Fig 2.**
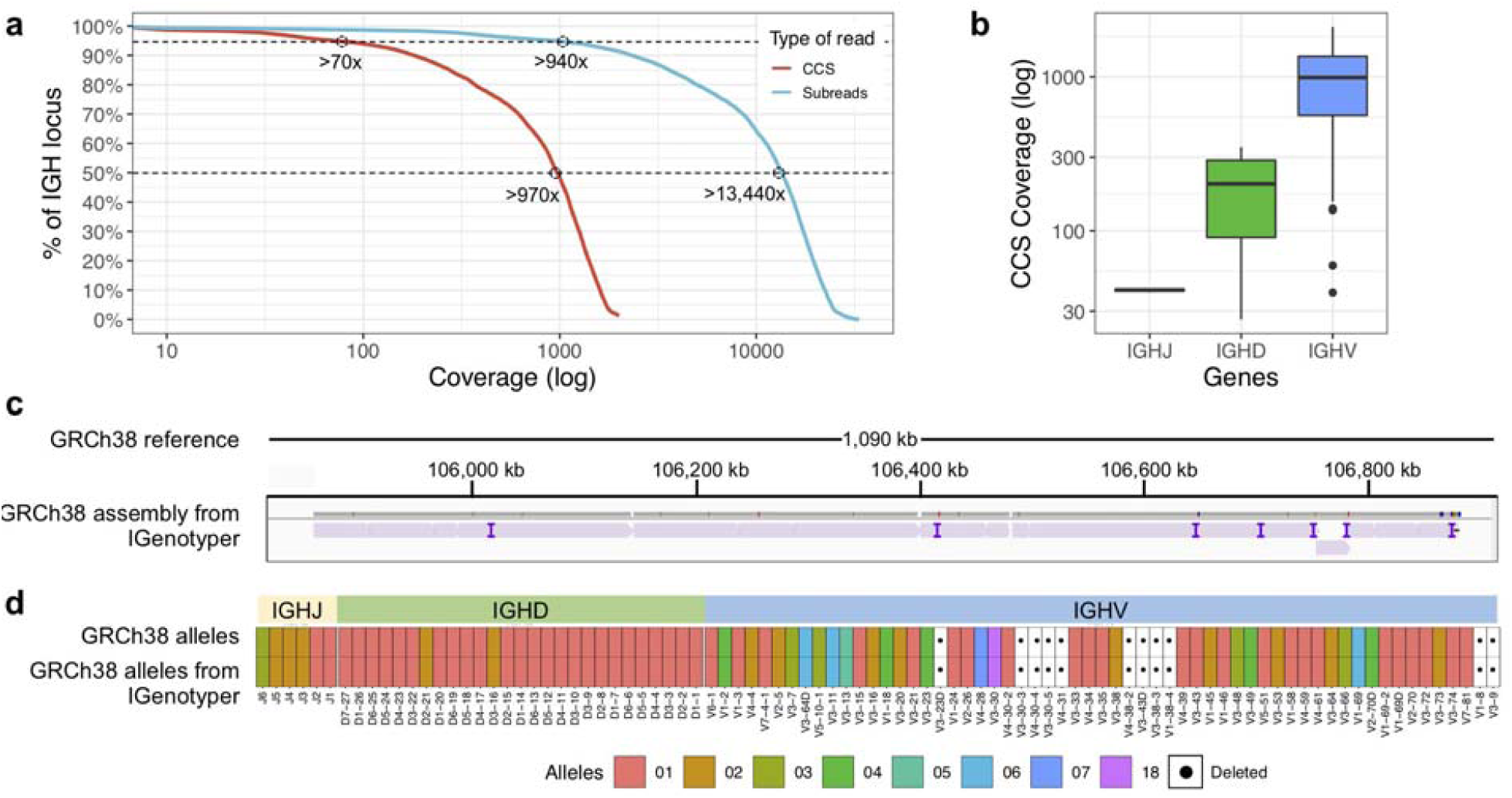
Benchmarking targeted long-read sequencing and assembly in a haploid DNA sample. **a** The empirical cumulative subread (blue) and CCS (red) coverage in IGH from the combined CHM1 dataset. The subread coverage for 95% and 50% (dotted line) of the locus is greater than 940x and 13,440x, and the CCS read coverage for 95% and 50% of the locus is greater than 70x and 970x, respectively. **b** CCS coverage across IGHJ, IGHD and IGHV genes. The average CCS coverage of IGHV genes was > 1000x. **c** IGenotyper assembly of CHM1 aligned to GRCh38. Purple tick marks represent indels in the IGenotyper assembly relative to GRCh38. **d** IGHJ, IGHD, and IGHV alleles detected by IGenotyper in CHM1 compared to alleles previously annotated in GRCh38.

All known SVs previously described in CHM1(*4*) were present in the IGenotyper assembly, accounting for all IGHV (n=47), IGHD (n=27), and IGHJ (n=6) F/ORF gene segments in this sample. In addition to genes previously characterized by BAC sequencing, the IGenotyper assembly additionally spanned *IGHV7-81*. Alleles identified by IGenotyper were 100% concordant with those identified previously in GRCh38 (Fig. 2d)(*4*).

### Assessing the accuracy of diploid IGH assemblies

We next assessed the ability of IGenotyper to resolve diploid assemblies in IGH, using a Yoruban (YRI; NA19240, NA19238, NA1239) and European (CEU; NA12878, NA12891, NA12892) trio from the 1000 Genomes Project(*51*) (1KGP; Fig. S3; Table S.4). Lymphoblastoid cell lines (LCLs), which are the primary source of 1KGP sample DNA are known to harbor V(D)J somatic rearrangements within the IG loci, including reduced coverage in IGHD, IGHJ, and proximal IGHV regions(*4, 18*). However, because IGenotyper assembles the IGH locus in a local haplotype-specific manner, V(D)J rearrangements can be easily detected (Fig. S4-5; Table S.7). Nonetheless, we focused our analysis exclusively on the IGHV region (9 Kb downstream of *IGHV6-1* to telomere) to avoid potential technical artifacts that would hinder our benchmarking assessment.

IGH enrichment was performed and libraries were sequenced on the RSII or Sequel platform (Table S.4). For diploid samples, IGenotyper (Fig. 1b) first identifies haplotype blocks using CCS reads that span multiple heterozygous SNVs within a sample. Within each haplotype block, CCS reads are then partitioned into their respective haplotype and assembled independently to derive assembly contigs representing each haplotype in that individual. Reads within blocks of homozygosity that cannot be phased are collectively assembled, as these blocks are considered to represent either: 1) homozygous regions, in which both haplotypes in the individual are presumed to be identical, or 2) hemizygous regions, in which the individual is presumed to harbor an insertion or deletion on only one chromosome (Fig. S6).

We assessed IGenotyper performance in the probands of each trio. IGenotyper assemblies were composed of 41 and 49 haplotype blocks in NA19240 and NA12878, respectively (Table S.8). Of these, 20/41 and 24/49 blocks in each respective sample were identified as heterozygous, in which haplotype-specific assemblies could be generated, totaling 826,548 bp (69.28%) in NA19240, and 424,834 bp (35.61%) in NA12878. Within these heterozygous blocks, the mean number of heterozygous positions was 76.16 (NA19240) and 52.08 (NA12878). Summing the bases assembled across both heterozygous and homozygous/hemizygous contigs, complete assemblies comprised 1.8 Mb of diploid resolved sequence in NA19240 and 1.4 Mb in NA12878 (Table 1). The difference in size is partially due to V(DJ) rearrangements and large deletions present in NA12878 relative to NA19240 (Fig. 3).

**Fig 3.**
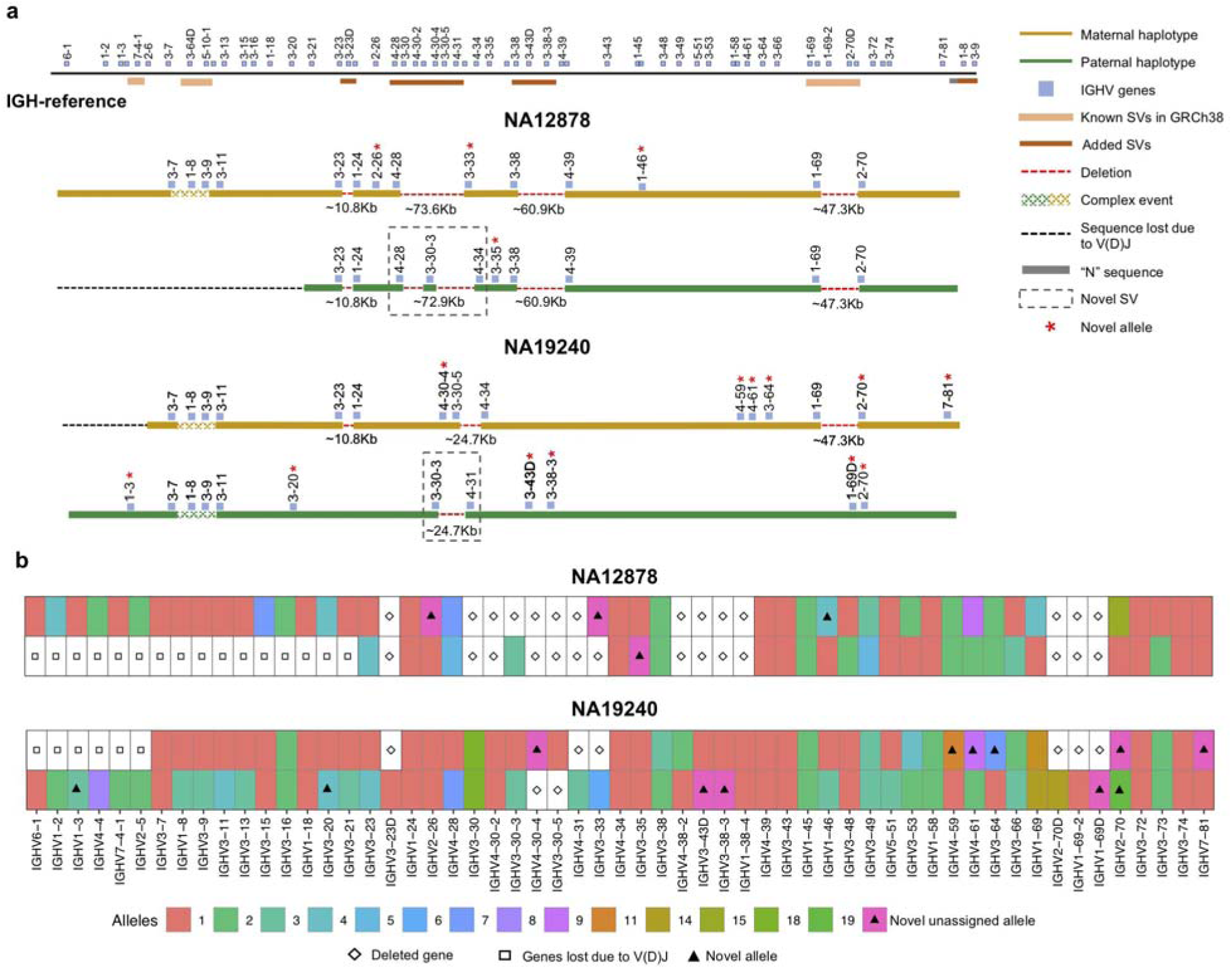
Haplotype-resolved assembly for characterizing structural variants and IGHV gene alleles in NA12878 and NA19240. **a** A schematic of the custom IGH-reference spanning the IGHV gene region (top). Brown bars indicate the positions of inserted SVs in the IGH-reference; pink bars indicate the positions of additional known SVs present in GRCh38 relative to GRCh37. Positions of IGHV (blue) genes are also indicated. Schematic depictions of resolved maternally (gold) and paternally (green) inherited haplotypes in NA12878 and NA19240 are shown. Detected deletions within annotated SV regions are labeled with a red dotted line, with detected sizes of each event also provided.

We next validated the accuracy of these assemblies using several orthogonal datasets: Sanger- and SMRT-sequenced fosmid clones, and paired-end Illumina data (Table 1). The Sanger-sequenced fosmids(*4*)(n=6, NA19240; n=2, NA12878) spanned 210.4 Kb of the NA19240 IGenotyper assembly and 70.2 Kb of the NA12878 assembly (Table S.9). The percent identity relative to the Sanger-sequenced fosmids was 99.989% for NA19240 and 100% NA12878. We also compared IGenotyper assemblies to additional fosmid assemblies in these samples (Rodriguez et al., *unpublished data*) sequenced using SMRT sequencing (n=85, NA19240; n=73, NA12878). These collectively spanned 1.5 Mb (82%; NA19240) and 1.2 Mb (82%; NA12878) of the IGenotyper assemblies, aligning with 99.996% and 99.995% sequence identity, respectively (Table 1). The numbers of putative indel errors were 276 (NA19240) and 188 (NA12878) (Table S.5). Of these indels, 254/276 and 180/188 were 1-2 bp indels, 71.26% (181/254) and 63.33% (114/180) of which were within homopolymers (Table S.6). Finally, we additionally assessed assembly accuracy using publicly available 30x PCR-free paired-end Illumina NovaSeq data from NA19240 and NA12878(*51*). We observed a total of 25 discordant bases and 143 indels in the NA19240 assembly (accuracy=99.989%), and 45 discordant bases and 154 indels in the NA12878 assembly (accuracy=99.991%; Table 1).

### Assessing local phasing accuracy and extending haplotype-specific assemblies with long-range phasing information

To assess local phasing accuracy, we also profiled the IGH locus in the parents of NA19240 and NA12878 (Table S.4). IGenotyper uses read-backed phasing to delineate reads within haplotype blocks prior to assembly. We tested the accuracy of local phasing (variant phasing within each haplotype block) by comparing read-backed and trio-based phased genotypes. No phase-switch errors were observed in the heterozygous haplotype blocks (n=20 blocks, NA19240; n=24, NA12878). Within homozygous blocks (excluding known SV sites), 27/57,313 (0.05%; NA12940) and 23/139,029 (0.02%; NA12878) bases did not follow a Mendelian inheritance pattern (Table S.10).

In both NA19240 and NA12878, we observe low localized read coverage in various regions of the locus, representing known technical limitations of probe-based DNA capture(*52*). Because of this, as well as regions of homozygosity/hemizygosity, IGenotyper is limited in its ability to generate phased haplotype assemblies across the entirety of the locus. However, we reasoned that with long-range phase information (e.g., trio genotypes) contigs from an IGenotyper assembly can be assigned to parental haplotypes. To assess this, heterozygous SNVs in NA19240 and NA12878 were phased using both sequencing reads and parental SNVs, resulting in completely phased haplotypes. To determine potential impacts on accuracy using either the local or long-range phasing approach, we compared each assembly type in the probands. Only 12 (NA12940) and 7 (NA12878) base differences were found between the locally phased and long-range phased assemblies. Taken together, these data suggest that individual contig assemblies generated by IGenotyper have high phasing accuracy.

We anticipate that alternative forms of long-range phasing data will be available in the future. One example would be IGHV, IGHD, and IGHJ haplotype information inferred from AIRR-seq data(*16, 17*). We assessed whether AIRR-seq based haplotype inference could be theoretically applied, by identifying the number haplotype blocks harboring heterozygous IGHV gene segments. In NA19240 and NA12878, 10/20 and 6/24 of the assembled heterozygous contig blocks harbored at least one heterozygous IGHV gene. In total, this equated to 53.5% (442,057 bps) in NA12940 and 80.72% (342,942 bps) in NA12878 of heterozygous bases that could theoretically be linked using this type of coarse long-range phase information, highlighting the potential strength of pairing the two approaches in larger numbers of samples.

### IGenotyper produces accurate gene annotation, SNV, indel, and SV variant call sets from diploid assemblies

Previous studies have demonstrated that assembling diploid genomes in a haplotype-specific manner increases the accuracy of variant detection(*37, 38, 42, 53–56*) and facilitates greater resolution on the full spectrum of variant classes(*57*). In addition to IGH locus assembly, IGenotyper detects SNVs, short indels, and SVs, including SNV calls within previously characterized complex SV/insertion regions. IGenotyper also provides direct genotypes for five previously described biallelic SVs (see Table S.11). This excludes the structurally complex *IGHV3-30* gene region, known to harbor multiple complex haplotypes; however, IGenotyper assemblies can be used for manual curation of this region (see below).

Using the fully phased diploid assemblies from each proband (Fig. 3), we assessed the validity of annotations/variant calls. We compared proband gene annotations and variant call sets to fosmid and parental assembly data (Table 2). In each sample, we noted the presence of a VDJ recombination event on one chromosome, which resulted in the artificial loss of alleles (Fig. 3a). However, because these events were detectable, they did not preclude our ability to make accurate annotations and variant calls.

**Table 2.**
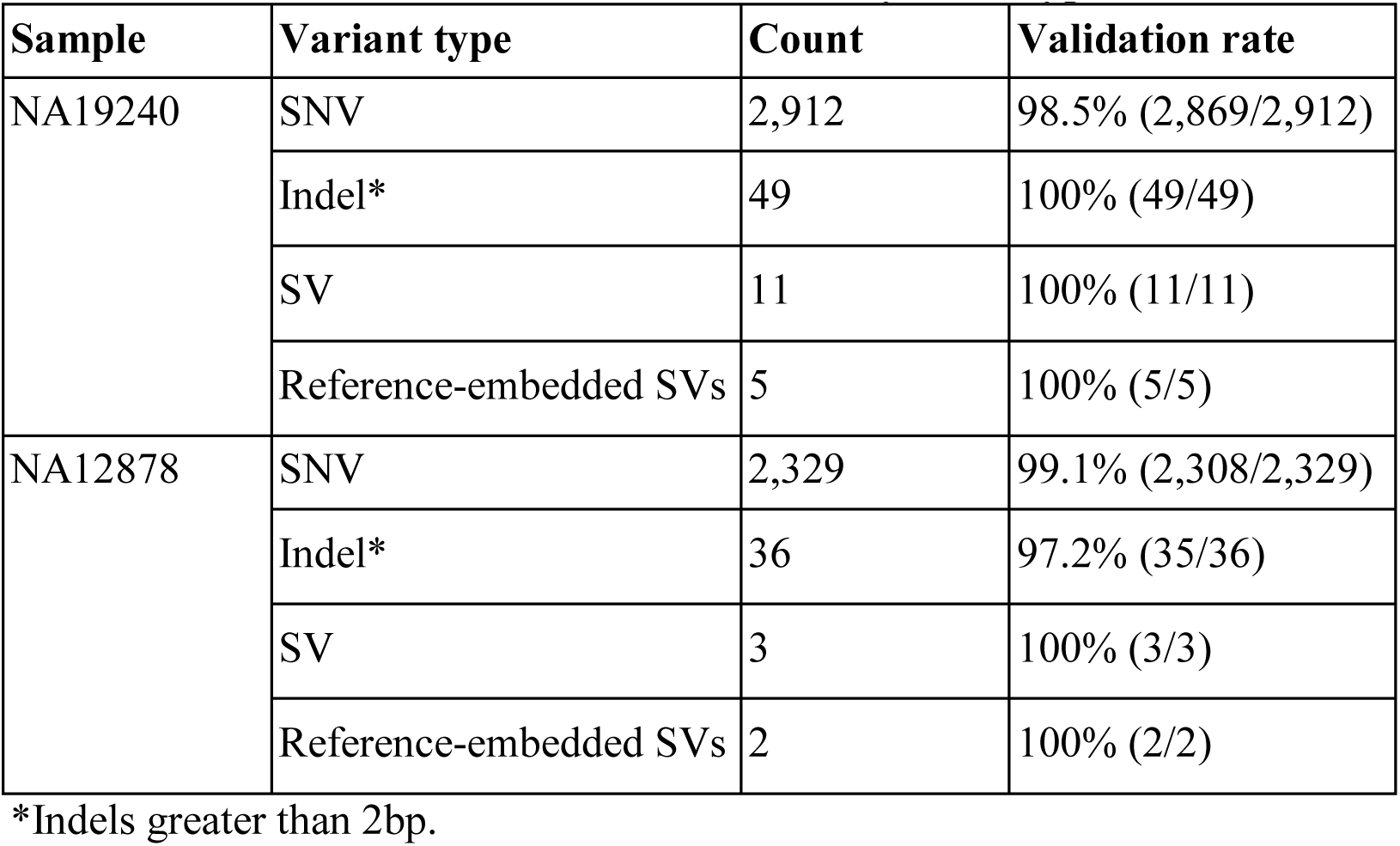
Count of different variants identified by IGenotyper

| Sample | Variant type | Count | Validation rate |
| --- | --- | --- | --- |
| NA19240 | SNV | 2,912 | 98.5% (2,869/2,912) |
|  | Indel* | 49 | 100% (49/49) |
|  | SV | 11 | 100% (11/11) |
|  | Reference-embedded SVs | 5 | 100% (5/5) |
| NA12878 | SNV | 2,329 | 99.1% (2,308/2,329) |
|  | Indel* | 36 | 97.2% (35/36) |
|  | SV | 3 | 100% (3/3) |
|  | Reference-embedded SVs | 2 | 100% (2/2) |

\*Indels greater than 2bp.

In NA19240, IGenotyper identified 79 unique non-redundant alleles across 57 IGHV genes (Fig. 3b; Table S.12, 13); 12 of these alleles were not found in IMGT, representing novel alleles. All 79 alleles were validated by parental and/or fosmid assembly data (Table S.12). In NA12878, 56 non-redundant alleles were called at 44 IGHV genes (Fig. 3b; Table S.14, 15), four of which were novel; all 56 alleles were validated (Table S.14).

Across IGHV we identified 2,912 SNVs, 49 indels (2-49 bps), and 11 SVs (> 50 bps) in NA19240. Collectively, IGenotyper-based genotypes for the parents of NA19240 and/or genotypes from the fosmids supported 2,869/2,912 SNVs, 31/36 indels, and 11/11 SVs in NA19240. In NA12878, we identified 2,329 SNVs, 36 indels (2-49 bps), and 3 SVs (> 50 bps), 2,308 (SNVs), 20 (indels), and 3 (SVs) of which were supported by orthogonal data. Included in the SVs called from both probands were events within previously identified SV regions (Fig. 1a; Table S.11). All of these regions are polymorphic at the population level(*4, 19, 31*), and several involve complex duplications and repeat structures (Fig. 3a). Strong concordance was observed in these regions between proband IGenotyper assemblies, fosmid clones, and parental CCS reads/assemblies (Fig. S7-12; Table S.16).

Additionally, we discovered novel SVs in both NA12878 and NA19240 within the region spanning the genes *IGHV4-28* to *IGHV4-34* (Fig. 3a). This site is a known hotspot of structural polymorphisms, in which six SV haplotypes have been fully or partially resolved(*4*). The longest haplotype characterized to date(*4*) contains four ∼25 Kb segmental duplication blocks. The novel SV haplotype in NA12878 contains a single ∼25 Kb segmental duplication block, and lacks 6 of the functional/ORF IGHV genes in this region. The novel SV haplotype in NA19240 contains 3/4 segmental duplication blocks, only lacking the genes *IGHV4-30-4* and *IGHV3-30-5*. Both of these novel SVs are supported by fosmid clones and parental data (Fig. S10).

### Identifying false-negative and -positive IGH variants in public datasets

Pitfalls of using short-read data for IGH variant detection and gene annotation have been discussed previously(*20, 58*). We assessed potential advantages of our approach compared to other high-throughput alternatives. In the CHM1 dataset, we defined a ground truth IGH SNV dataset by directly aligning the IGH locus haplotype from GRCh38(*4*) to that of GRCh37(*50*). We identified 2,940 SNVs between the two haplotypes in regions of overlap (i.e., non-SV regions). We next aligned Illumina paired-end sequencing data generated from CHM1(*38*) and our CHM1 IGenotyper assemblies to GRCh37. We detected 4,433 IGH SNVs in the Illumina dataset, and 2,958 SNVs in the IGenotyper assembly. The Illumina call set included only 73.2% (2,153) of the ground truth SNVs, as well as an additional 2,274 false-positive SNVs (Fig. 4a). In contrast, the IGenotyper call set included 99.0% (2,912) of the ground truth SNVs, and only 46 (1.6%) false-positive SNVs were called (Fig. 4a).

**Fig 4.**
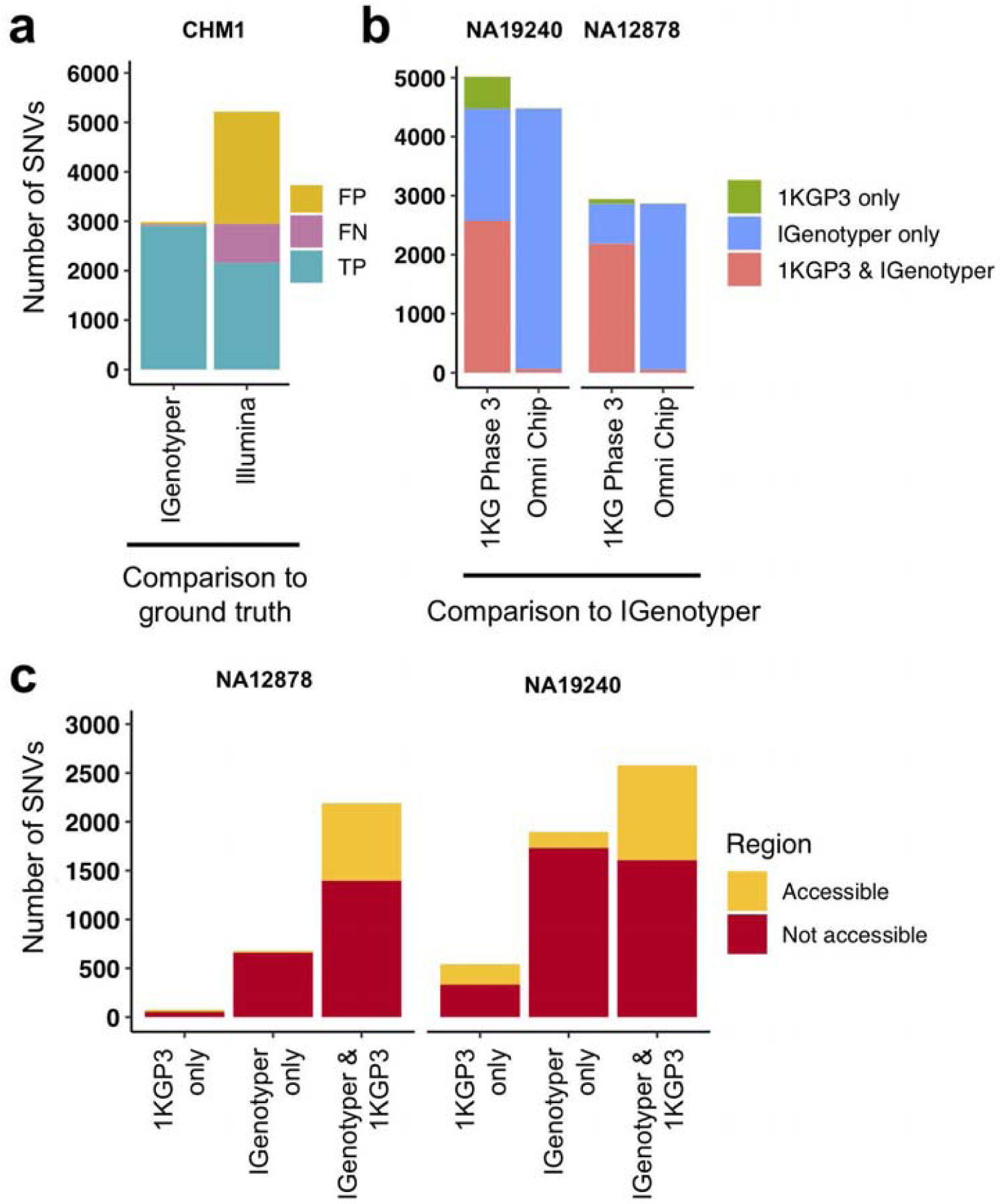
Comparison of SNVs identified by IGenotyper to SNVs called using short-read and microarray data. Genes directly flanking or within detected SVs are labelled. Genes with novel alleles are labelled with red asterisks, and novel SVs are indicated with dotted boxes. **b** Alleles predicted by IGenotyper for each gene across both haplotypes in NA12878 and NA19240. Novel alleles are marked by a filled triangle and deleted genes and genes not present due to V(D)J recombination are marked by a square and diamond, respectively.

We next compared IGenotyper genotypes for NA19240 and NA12878 to 1KGP short-read and microarray data (Fig. 4b). IGenotyper SNVs were lifted over to GRCh37 (n=4,474, NA19240; n=2,868, NA12878), excluding SNVs within SV regions not present in GRCh37 (n=703, NA19240; n=737, NA12878) Table S.17). In total, only 57.6% (2,578/4,474) and 76.4% (2,190/2,868) of the IGenotyper SNVs were present in the 1KGP call set for NA19240 and NA12878 (Fig. 4b; Table S.18). Critically, because insertion SVs are not present in GRCh37, the additional SV-associated SNVs were also missed. Thus, in total, 50.2% (2,599/5,177) and 39.3% (1,415/3,605) of IGenotyper SNVs were absent from 1KGP (Fig. 4b), including SNVs within 18 and 6 IGHV genes; 1,350/4,474 (NA19240) and 526/2,868 (NA12878) IGenotyper SNVs were not found in any 1KGP sample. The 1KGP call set also included an additional 542 (17.4%) and 76 (3.4%) SNVs (putative false-positives) for NA19240 and NA12878, respectively, including putative false-positive SNVs in 6 and 3 IGHV genes (Fig. 4b). In contrast to SNVs found only in the 1KGP datasets, we found that in both probands >90% of SNVs detected only by IGenotyper were within NGS-inaccessible regions (NA19240, 91.3%, 1,731/1,896; NA12878, 97.1%, 658/678; Fig. 4c; Table S.19), suggesting that IGenotyper offers improvements in regions that are inherently problematic for short-reads. We additionally assessed Hardy-Weinberg Equilibrium (HWE) at the interrogated SNVs, as deviation from HWE is often used to assess SNV quality. In both probands, we found that a greater proportion of SNVs unique to the 1KGP callset deviated from HWE than those called by IGenotyper (Fig. S13).

Finally, we compared larger variants, indels and SVs identified by IGenotyper to those detected by the Human Genome Structural Variation Consortium (HGSV) in NA19240(*37*). First we assessed support for six large known identified SVs in NA19240 (Fig. 3a; Tables S11 and S21). BioNano data detected events in five out of the six SV regions. The complex SV spanning *IGHV1-8*/*3-9*/*IGHV3-64D*/*5-10-1* was not detected, likely because this event involves a swap of sequences of similar size (∼38 Kb)(*4*), making it difficult to identify using BioNano. A critical difference to BioNano is that IGenotyper provides nucleotide level resolution allowing for fuller characterization of SV sequence content, including SNVs within these regions (as noted above). In addition to these large SVs, IGenotyper also identified 39 indels (3 - 49 bps) and an additional 11 SVs (>49 bps; 57-428 bps) in NA19240. Of these, 20 indels and 9 SVs were present in the HGSV integration call set.

### Effects of false-positive and -negative variants on imputation accuracy

We explored the potential advantage of our genotyping approach compared to array genotyping and imputation. We applied our long-read capture method to a sample selected from a recent rheumatic heart disease (RHD) GWAS(*22*), which identified IGH as the primary risk locus. Direct genotyping in this sample was carried out previously using the HumanCore-24 BeadChip (n=14 SNVs) and targeted Sanger sequencing (n=8 SNVs); genotypes at additional SNVs were imputed with IMPUTE2(*59*), using a combination of 1KGP and population-specific sequencing data as a reference set. We compared IGenotyper SNVs from this sample to directly genotyped variants and imputed variants selected at three hard call thresholds (0.01, 0.05, and 0.1; Fig. 5). The majority of directly genotyped SNVs (array, 13/14; Sanger, 8/8) were validated by IGenotyper. However, the validation rate varied considerably for the imputed SNVs, depending on the hard call threshold used (Fig. 5a; 93.8%, 0.01; 92.7%, 0.05; 40.5%, 0.1), with the signal to noise ratio decreasing significantly from the 0.01 to the 0.1 threshold (Fig. 5b). In all cases, IGenotyper called a substantial number of additional SNVs (Fig. 5a); the majority of these were located in the proximal region of the locus, which was poorly represented by both directly genotyped and imputed SNVs (Fig. 5c).

**Fig 5.**
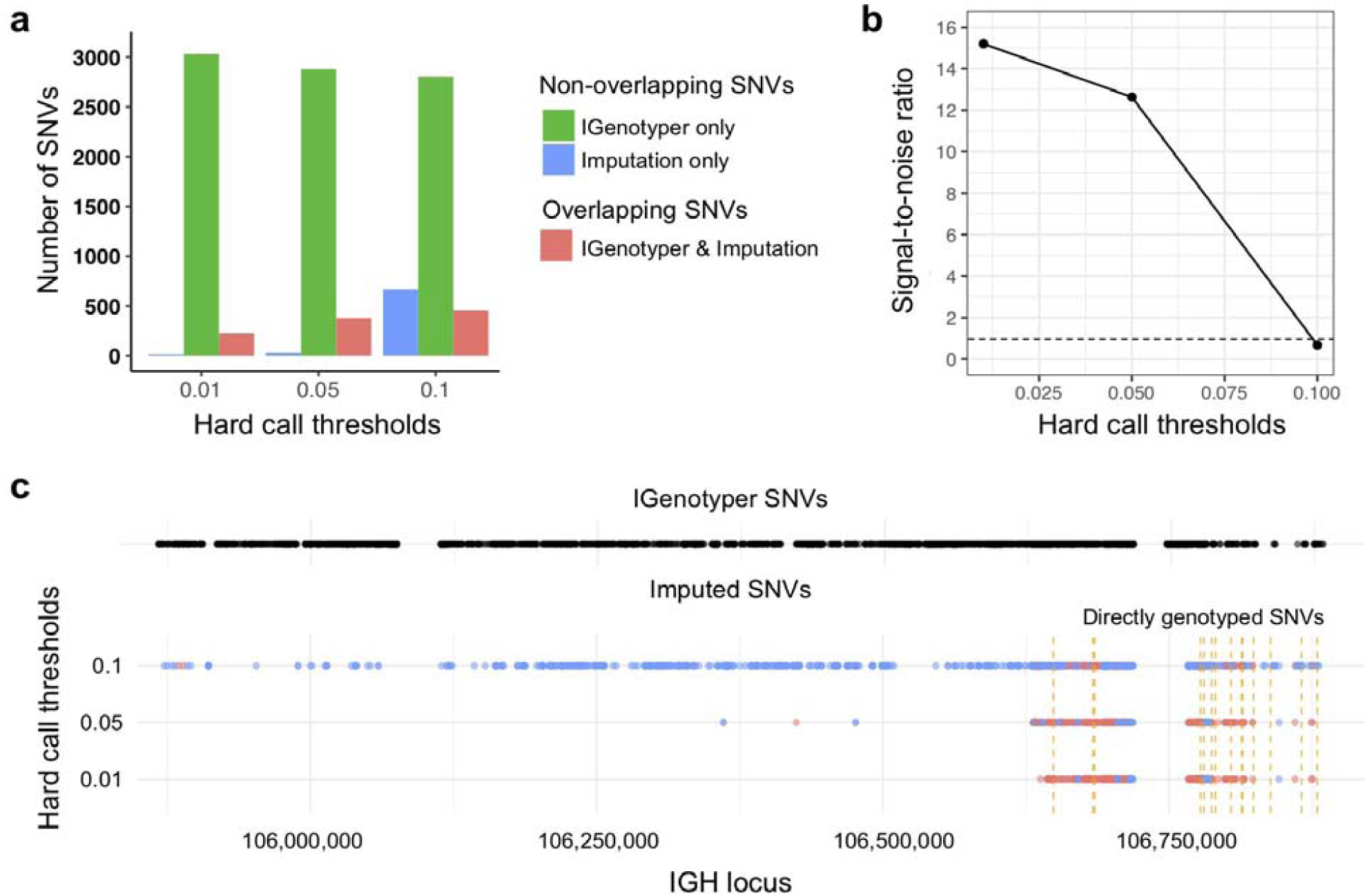
Comparison of IGenotyper variant calls to microarray-/imputation-based genotyping. **a** SNVs detected by IGenotyper were compared to array-based and imputed SNVs in a sample from a recent RHD GWAS. Imputed SNVs were filtered based on different hard call thresholds (0.01, 0.05, and 0.1). Each set of filtered SNVs was compared to IGenotyper SNVs. The number of imputed SNVs not identified by IGenotyper (blue), SNVs only detected by IGenotyper (green), and overlapping SNVs (red) are shown above. **b** The signal-to-noise ratio was calculated using the overlapping SNVs between IGenotyper and imputation, and the SNVs identified only by imputation (not called by IGenotyper). Below the signal-to-noise ratio of 1 (dotted line), there is more noise than signal. **c** Positions of SNVs detected by IGenotyper and at different hard call thresholds. Each point represents a SNV. Blue SNVs are SNVs only identified by imputation and not detected by IGenotyper, and red SNVs are SNVs found both by imputation and IGenotyper. **a** SNVs detected by IGenotyper and Illumina/GATK in CHM1 were compared to a CHM1 ground truth SNV dataset; numbers of false-negative, false-positive, and true-positive SNVs in each callset are shown. **b** SNVs in the 1KGP Phase 3 datasets were compared to SNVs detected by IGenotyper in NA19240 and NA12878. The total number of SNVs in each bar sums to the number of overlapping SNVs and the number of SNVs unique to each dataset. **c** SNVs found by IGenotyper and the 1KGP3 dataset, found only by IGenotyper and found only in the 1KGP3 dataset were partitioned into regions identified as accessible by the 1KGP accessible genome browser track.

### Sample multiplexing leads to reproducible assemblies and variant calls

An advantage to the capture-based approach employed here is the ability to multiplex samples. To demonstrate this, eight technical replicates of NA12878 were captured, barcoded, pooled, and sequenced on a single Sequel SMRT cell (Table S.22), yielding a mean CCS coverage ranging from 41.3-101.6x for each library (Fig. 6a). We then simulated different plexes (2-, 4-, 16-, 24-, 40-plex) by either combining or partitioning data from this 8 plex, allowing us to assess the impacts of read depth on IGH locus coverage, assembly accuracy, and variant calling. The mean CCS coverage per plex ranged from 308.7X (2-plex) to 15.5X (40-plex) (Fig. 6a). To compare IGenotyper metrics across plexes, we chose the 2-plex sample with the highest CCS coverage to use as the ground-truth dataset; all other assemblies and variant call sets were compared to this sample. The lower coverage 2-plex assembly covered 99.75% of the ground truth assembly with a sequence identity concordance of 99.99%. For the remaining comparisons, mean assembly coverage and sequence concordance estimates ranged from 99.27% (4-plex) to 86.95% (40-plex), and 99.99% (4-plex) to 99.99% (40-plex). The mean number of observed SNVs ranged from 2,471 (2-plex) to 1,936 (40-plex) (Fig. 6b). When comparing these to ground truth SNVs, we observed high recall rates (>80%), even among the 40-plex assemblies (Fig. 6c); recall was >90% for all but one of the 4- and 8-plex assemblies (Fig. 6c). Importantly, although the recall rate of true-positive SNPs decreased as expected in higher plexes, we observed very little variation in the false-positive rate (Fig. 6c).

**Fig 6.**
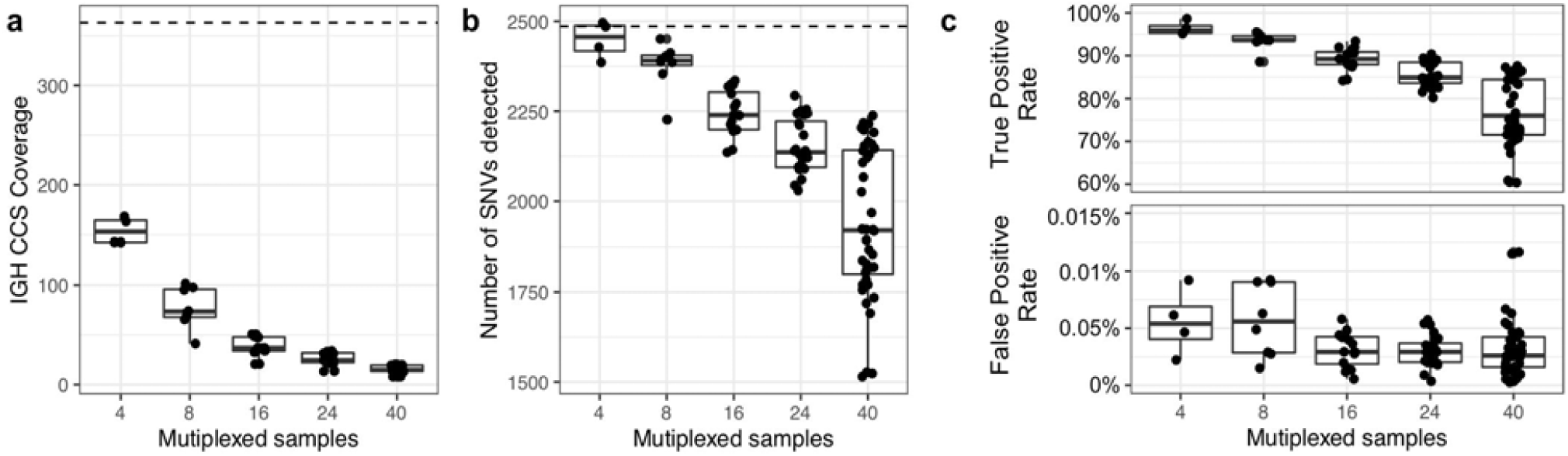
Assessment of sample multiplexing on assembly coverage and variant calling. Eight replicates of NA12878 were multiplexed and sequenced on a single Sequel SMRT cell. The eight replicates were combined or down sampled to simulate a sequencing run with 2, 4, 16, 24 and 40 samples. The simulated sample from the 2-plex run with the highest coverage was treated as the ground truth. **a** CCS coverage of sequencing runs with different numbers of multiplexed samples. The dotted line represents the coverage of a sample from a 2-plex sequencing run. **b** Number of SNVs found across samples per each multiplexed sequencing run. The dotted line represents the number of SNVs detected in a sample from a 2-plex sequencing run. **c** True and false positive rate of SNVs of each sample in each multiplexed sequencing run. The SNVs from each sample were compared to SNVs from a sample sequenced in a 2-plex run.

## Discussion

For decades, comprehensive genetic analysis of the human IGH locus has been intractable(*20, 21, 58*). As a result, our understanding of the extent of IGH germline diversity in human populations, and how this diversity contributes to B cell mediated immunity remains incomplete(*20, 21*). Here, we have leveraged existing genomic haplotype data to design a novel IGH assembly and genotyping framework that combines targeted long-read sequencing with a novel bioinformatics toolkit (IGenotyper). This end-to-end pipeline can reconstruct completely phased assemblies via the integration of long-range phase information. Utilizing high-quality CCS reads and derived assemblies facilitates characterization of IGH gene segments and all forms of coding and non-coding variants, including the discovery of novel variants and IG alleles.

We validated our pipeline on eight ethnically diverse human samples with orthogonal data that highlighted multiple strengths of our approach. First, we chose individuals with available large-insert, clone-based assembly datasets for direct comparisons to capture/IGenotyper assemblies, as well as Illumina short-read data for assessing assembly error correction metrics. These comparisons revealed high concordance between assemblies (>99%) in both haploid and diploid samples. Second, we directly validated variants and alleles using trio data. Finally, comparisons to additional variant call sets (1KGP and HGSV) allowed us to assess concordance in variant detection (including indels and SVs), and demonstrate advantages over alternative high-throughput methods. Specifically, compared to microarray-based and short-read sequencing methods, IGenotyper variant call sets were more comprehensive, exhibited greater locus coverage and were more accurate.

Much recent effort has focused on identifying IG genes/alleles absent from existing databases(*3–7, 10, 18, 60–63*), revealing many undiscovered alleles in the human population. In our analysis, we identified 16 novel alleles from only two samples. Consistent with previous suggestions of undersampling in non-Caucasian populations(*6*), the majority (n=12) were in the Yoruban individual, for which 6 additional novel alleles had been reported in an earlier study(*4, 50*). Notably, the novel alleles described in NA19240 represent the largest contribution to the IMGT database from a single individual. Related to this point, we also observed high numbers of false-positive/negative IGHV SNVs in 1KGP datasets, reinforcing that efforts to identify IG alleles from 1KGP data should be done with extreme caution(*58, 64, 65*). An added advantage of our approach is the ability to capture variation outside of IG coding segments and more fully characterize SVs. Although several studies have begun to demonstrate the extent of SV haplotype variation(*4, 16, 17, 63*), information on polymorphisms within these SVs, and within IGH regulatory and intergenic space remains sparse(*21*). It is worth noting that, in the few samples analyzed here, the majority of variants were detected in non-coding regions, including SNVs within RS, leader, intronic, and intergenic sequences. We also showed that IGenotyper resolved novel SVs within the complex *IGHV3-30* gene region in both 1KGP diploid samples. Together, these examples are testament to the fact that our approach represents a powerful tool for characterizing novel IGH variation.

Despite evidence that IG polymorphism impacts inter-individual variation in the antibody response(*12, 19, 24, 29*), the role of germline variation in antibody function and disease has not been thoroughly investigated. The population-scale IGH screening that will be enabled by this approach will be critical for conducting eQTL studies and integrating additional functional genomic data types to better resolve mechanisms underlying IG locus regulation, which have only so far been applied effectively in model organisms(*66–69*). Delineating these connections between IGH polymorphism and Ab regulation and function will be critical for understanding genetic contributions to Ab mediated clinical phenotypes(*21*).

To date, few diseases have been robustly associated to IGH(*22, 23, 70, 71*). We previously suggested this was due to sparse locus coverage of genotyping arrays and an inability of array SNPs to tag functional IGH variants(*4, 20*). We have provided further support for this idea here. First, the identification of both putative false negative and positive SNVs in 1KGP samples highlights potential issues with imputation-based approaches using 1KGP samples as a reference set. Second, our direct analysis of capture/IGenotyper data in a sample from a recently conducted GWAS(*22*) also demonstrated that IGenotyper resulted in a larger set of genotypes, with improved locus coverage compared to imputation. Together, these analyses highlight the potential for our framework to supplement GWASs for both discovery and fine mapping efforts, and through building more robust imputation panels. A strength of our approach is that the user can determine the sequencing depth and locus coverage, depending on whether the intent is to conduct full-locus assemblies or genotyping screens; although the number of detected variants decreases with increased multiplexing, false-positive rates remain low. The recently released, higher throughput Sequel 2 platform, in combination with read length improvements, will allow for processing of larger cohorts at low cost.

A current technical limitation of our framework is the decreased efficiency of probes in particular regions of IGH. However, we showed that these regions represent a small fraction of IGH, with negligible impacts on locus coverage and assembly quality. Future iterations of target-enrichment protocols will improve efficiency through methodological or reagent modifications. A key strength of IGenotyper is its flexibility to accommodate other data types; e.g., users interested in complete haplotype characterization can already provide long-range phase information to inform a complete diploid assembly. We envision other forms of data will be leveraged in future applications. A second limitation is the potential to miss unknown sequences not specifically targeted by capture probes. We expect this issue to be mitigated in the future as more IG haplotypes are sequenced using our method, large-insert clones, or WGS approaches; enumeration of these data will allow for refinement on protocol design and IGenotyper functionality. Ultimately, we promote an advance toward a more extensive collection of IGH haplotype reference datasets and variants as a means to leverage a population reference graph approach, which has been shown to be effective in other hyperpolymorphic immune loci(*72*).

To the best of our knowledge, this is the first combined molecular protocol and analytical pipeline that can provide comprehensive genotype and annotation information across the IGH locus, with the added ability to be applied to a large number of samples in a high-throughput manner. Given the importance of antibody repertoire profiling in health and disease, characterizing germline variation in the IG regions will continue to become increasingly important. Our strategy moves towards the complete ascertainment of IG germline variation, a prerequisite for understanding the genetic basis of Ab-mediated processes in human disease(*73*).

## Materials and Methods

### Library preparation and sequencing

Genomic DNA samples (1K Genomes DNA) were procured from Coriell Repositories (Camden, NJ) and collaborators (CHM1). Briefly, two micrograms of high molecular weight genomic DNA from each sample was sheared using g0-tube to ∼8 Kb (Covaris, Woburn, MA). These sheared gDNA samples were size selected to include 5 Kb-9 Kb fragments with Blue Pippin (Sage Science, Beverly MA). Following size selection, each sample was End Repaired and A-tailed following the standard KAPA library preparation protocol (Roche, Basel, Switzerland). For multiplexed samples, adapters containing sequence barcodes (Pacific Biosciences, Menlo Park, CA) and a universal priming sequence were ligated onto each sample. Each sample was PCR amplified for 9 cycles using HS LA Taq (Takara, Mountain View, CA) and cleaned with 0.7X AMPure beads to remove small fragments and excess reagents (Beckman Coulter, Brea, CA). The genomic DNA libraries were then captured following the SeqCap protocol which was modified to increase final capture reaction volume by 1.5X (Roche, Basel, Switzerland).

Following capture, the libraries were washed following SeqCap protocol, substituting vortexing with gentle flicking. The washed capture libraries were PCR amplified for 18 cycles using HS LA Taq and cleaned with 0.7X AMPure beads.

Capture libraries were prepared for PacBio sequencing using the SMRTbell Template Preparation Kit 1.0 (Pacific Biosciences, Menlo Park, CA). Briefly, each sample was treated with a DNA Damage Repair and End Repair mix in order to repair nicked DNA and add A-tails to blunt ends. SMRTbell adapters were ligated onto each capture library to complete SMRTbell construction. The SMRTbell libraries were then treated with exonuclease III and VII to remove any unligated gDNA and cleaned with 0.45X AMPure PB beads (Pacific Biosciences, Menlo Park, CA). Resulting libraries were prepared for sequencing according to protocol and sequenced as single libraries per SMRTcell with P6/C4 chemistry and 6h movies on the RSII system, or as multiplexed libraries per SMRTcell 1M, annealed to primer V4 and sequenced using 3.0 chemistry and 20h movies, on the Sequel system (see Table S.4 for details).

### Creating a custom IGH locus reference

The IGH locus (chr14:106,326,710-107,349,540) was removed from the human genome reference build GRCh37, and the expanded custom IGH locus reference was inserted in its place. The expanded custom IGH locus is the GRCh38 IGH locus with extra IGH sequences inserted. The extra IGH sequences include sequence from fosmid clones AC244473.3, AC241995, AC234225, AC233755, KC162926, KC162924, AC231260, AC244456 and KC162925, and sequence from human genome reference build GRCh37 IGH locus (chr14:106,527,905-106,568,151). The IGH sequence from fosmid clones are sequences not found in the GRCh38 IGH locus. The IGenotyper toolkit command ‘IG-make-ref’ takes as input the human genome reference build GRCh37 and creates the custom IGH locus reference.

### IGenotyper: a streamlined analysis tool for IGH locus assembly, variant detection/genotyping, and gene feature annotation

Running IGenotyper returns multiple output files: (1) the alignment of the CCS reads and assembled locus to the reference in BAM format; (2) the assembled IGH locus in FASTA format, (3) the SNVs in VCF; (4) indels and SVs in BED format; (5) a parsable file with genotyped SVs; (6) a parsable file with the detected alleles for each functional/ORF gene; and several tab delimited files detailing different sequencing run and assembly statistics. The BAM file contains phased CCS reads and includes haplotype annotation in the read group tag of every read. This allows the user to separate the reads into their respective haplotype in the Integrative Genomics Viewer (IGV) visualization tool(*74*). The VCF file contains annotations indicating whether SNVs reside within SV regions, and IG gene features, including coding, intron, leader part 1 (LP1), and recombination signal (RS) sequences. A user-friendly summary file is produced with links to output files (see Fig. 7; sample summary output for NA19240), including summary tables and figures pertaining to: locus sequence coverage; counts of SNVs, indels and SVs; allele annotations/genotypes for each IGHV, IGHD, and IGHJ genes; and lists of novel alleles.

**Fig. 7.**
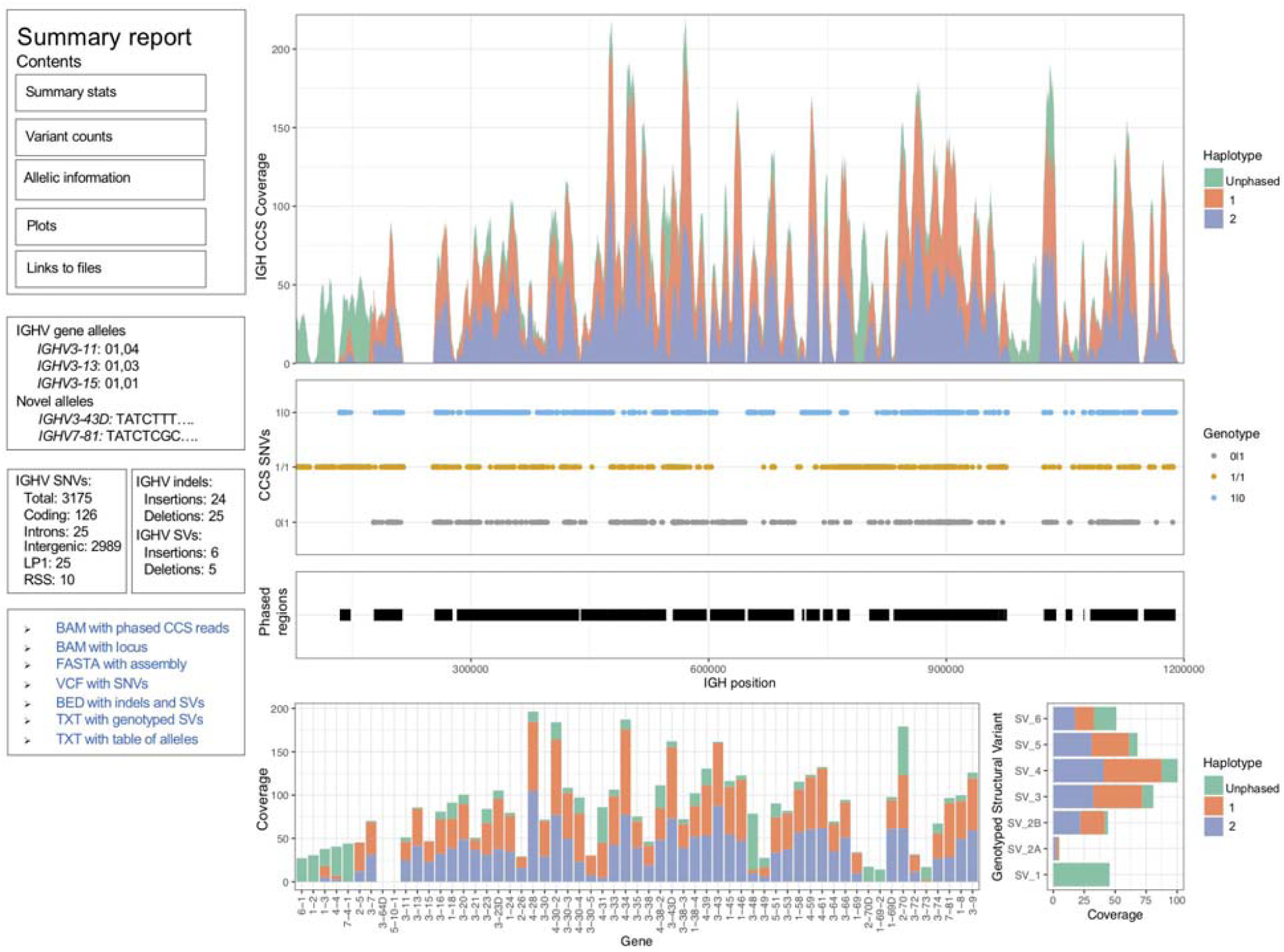
HTML summary report from IGenotyper. IGenotyper produces a summary report with tables containing different sequencing and assembly summary statistics, the count of variants (SNVs, indels, and SVs), genotypes of the IGHV,D, and J genes and a link to files with the variants and alleles (VCF, BED and TXT files).

### Running IGenotyper

IGenotyper has three main commands: ‘phase’, ‘assemble’, and ‘detect’. The input is the subread bam output from the RS2 or Sequel sequencing run. ‘phase’ phases the subreads and CCS reads. ‘assemble’ partitions the IGH locus into haplotype-specific regions and assembles each region. ‘detect’ detects SNVs, indels and SVs, genotypes 5 SVs embedded in the IGH reference and assigns the IGH genes to alleles from the IGMT database.

### Phasing SMRT sequencing reads

Raw SMRT sequences with at least 2 subreads are converted into consensus sequences using the ‘ccs’ command (https://github.com/PacificBiosciences/ccs). CCS reads and subreads are aligned to our expanded custom IGH locus reference using BLASR(*75*). Phased SNVs are detected from the CCS reads using the WhatsHap(*76*) ‘find_snv_candidates’, ‘genotype’, and ‘phase’ commands. The subreads and CCS reads are phased using the command ‘phase-bam’ from MsPAC(*54*).

### Assembling the IGH locus

Haplotype blocks are defined using the WhatsHap ‘stats’ command. Haplotype-specific CCS reads within each haplotype block are assembled separately using Canu(*77*); regions outside haplotype blocks are assembled using all aligned CCS reads. Regions lacking contigs recruit raw subreads and repeat the assembly process. Contigs with a quality score less than 20 are filtered.

### Detecting variants from SNVs to SVs

SNVs variants are detected from the assembly aligned to the reference. SNVs in the VCF are annotated with:

1. Contig id used to detect the SNV
2. Overlapping SV id (if any)
3. A true or false value if the SNV is also detected by the CCS reads
4. Whether the SNV falls within an intronic, leader part 1 or gene
5. Whether the SNV is within the IGHV, IGHD, or IGHJ region
6. For phased blocks, the haplotype block id and genotype

Indels and SVs are detected using the ‘sv-calling’ command from MsPAC(*54*). Importantly, each indel and SV is sequence-resolved since they are identified from a multiple sequence alignment using the haplotype-specific assemblies and reference.

### Assigning alleles to IGH genes extracted from the assembly

Assembled sequences overlapping the IGH genes are extracted and compared to the alleles in the IMGT database (v201938-4). Sequences not observed in the database are labelled as novel. CCS reads overlapping IGH gene sequences are also extracted and compared to the IMGT database. To provide supporting evidence for the allele, a count for the number of CCS reads with the same sequence found in the assembly is reported(currently only supported for Sequel data). Assembly and CCS sequences are outputted in a FASTA file; gene names, alleles (and allele sequence if novel), andread support are outputted in a tab-delimited file.

### Assessing accuracy of NA12878 and NA19240 assembly

The accuracy of the assembly of the NA19240 sample was assessed by BLAST (v2.7.1+) alignment of contigs to fosmids assembled from Sanger sequencing (NA19240 accession numbers: AC241513.1, AC234301.2, KC162926.1, AC233755.2, AC234135.3, AC244463.2; NA12878 accession numbers: AC245090.1, AC244490.2).

The accuracy of the assemblies was also accessed using Illumina data. Illumina HiSeq 2500 2×126 paired-end, PCR-free sequencing data (ERR894723, ERR894724, ERR899709, ERR899710 and ERR899711) was downloaded from the European Nucleotide Archive and aligned to the NA19240 assembly using bwa mem (v0.7.15-r1140). The total coverage across all sequencing runs was 75.6X and Pilon (v1.23) reported an accuracy of 99.996 (102 errors in 2,396,307 bp). Additionally, Illumina TruSeq 2×151 paired-end, PCR-free sequencing run ERR3239334 was downloaded from the European Nucleotide Archive and aligned to the NA12878 assembly. The coverage was 35.5X and Pilon reported an accuracy of 99.991 (167 errors in 1,918,794 bp).

### Manual curation of IGHV3-30 and IGHV1-69 gene regions

*IGHV3-30* and *IGHV1-69* gene duplication regions did not completely assemble into a single contig per haplotype, but instead were split into multiple contigs. To resolve these regions an additional curation step was employed: contigs were aligned to each other using BLAST and overlapping contigs with high alignment score were merged.

In NA19240, the *IGHV3-30* gene region duplication was initially assembled into 8 contigs. Two contigs were merged to form a novel SV containing a ∼25 Kb deletion relative to the IGH-reference. The two contigs overlapped by 7,706 bp with 5 bp mismatches and 9 gaps (11 gap bases). The alternate haplotype was initially assembled into 6 contigs. The 6 contigs overlapped by more than 2.3 Kbp with 0 bp mismatches and a total of 8 gap bases, allowing them to be merged into a single contig. Both haplotypes were validated with fosmids and assemblies from the parents (Supplementary Figure 10). The resulting contigs completely resolved the SVs on both haplotypes. This process was repeated for NA12878, and in both probands for the *IGHV1-69* gene region.

Leveraging parental and fosmid assembly data, we determined that NA19240 carried three distinct haplotypes within the SV region spanning *IGHV1-69, IGHV2-70D, IGHV1-69-2, IGHV1-69D*, and *IGHV2-70* (Fig. S12). An insertion haplotype carrying all genes within the region was paternally inherited; a deletion haplotype, lacking *IGHV2-70D, IGHV1-69-2*, and *IGHV1-69D*, was inherited from the mother; and a second deletion haplotype was detected in both the capture/IGenotyper and fosmid assembly data, but was not supported by either parental dataset. This deletion haplotype was identical to the paternally derived insertion haplotype on the flanks of the deletion event, suggesting it represented a somatic SV. Whether this arose natively in NA19240 or is an artifact found only within the LCL is not known. To construct the most accurate assemblies of the inherited haplotypes, we attempted to remove reads representing this somatic deletion and performed a local reassembly. This allowed us to produce more accurate contigs across this region which exhibited higher concordance to both fosmid and parental datasets (Fig. S12).

### Comparing SNVs from Illumina and IGenotyper on CHM1 sample

SNVs from our assembly and Illumina reference alignment to GRCh37 were compared to a ground truth SNV dataset generated by aligning GRCh38 region chr14:106,329,408-107,288,965 to GRCh37 using pbmm2 (v1.0.0). Positions with base differences in the alignment were labelled as SNVs. 2×126 TruSeq PCR-free Illumina libraries (SRR3099549 and SRR2842672) were aligned to GRCh37 with bwa (0.7.15-r1140) and SNVs were detected using the standard protocol with GATK (v3.6) tools, HaplotypeCaller and GenotypeGVCFs(*78*). The SNVs were filtered using bcftools (v1.9) for SNVs with genotype quality greater than 60 and read depth greater than 10.

### Comparing 1000 Genomes project SNVs to NA12878 and NA19240 SNVs detected by IGenotyper

SNVs detected by IGenotyper were compared to SNVs in the 1000 Genome Phase 3 project. IGH-specific SNVs from the 1000 Genomes Phase 3 project (ftp://ftp.1000genomes.ebi.ac.uk/vol1/ftp/release/20130502/supporting/hd_genotype_chip/ALL.chip.omni_broad_sanger_combined.20140818.snps.genotypes.vcf.gz, ftp://ftp.1000genomes.ebi.ac.uk/vol1/ftp/release/20130502/ALL.chr14.phase3_shapeit2_mvncall_integrated_v5a.20130502.genotypes.vcf.gz, ftp://ftp.1000genomes.ebi.ac.uk/vol1/ftp/release/20130502/supporting/hd_genotype_chip/ALL.chip.omni_broad_sanger_combined.20140818.snps.genotypes.vcf.gz, and ftp://ftp.1000genomes.ebi.ac.uk/vol1/ftp/phase3/data/NA19240/cg_data/NA19240_lcl_SRR832874.wgs.COMPLETE_GENOMICS.20130401.snps_indels_svs_meis.high_coverage.genotypes.vcf.gz) were extracted using ‘bcftools view --output-type v, --regions 14:106405609-107349540, - -min-ac 1, --types snps’. Overlap between the 1000 Genomes Phase 3 SNVs and SNVs detected by IGenotyper was determined using BEDTools ‘intersect’ command. Overlapping SNVs with discordant genotypes were labelled as discordant/non-overlapping SNVs.

### Comparing indels, SVs, and BioNano data from the 1000 Genome Structural Variation Consortium

Indels and SVs detected by IGenotyper were compared to indels and SVs from the Human Genome Structural Variation (HGSV) Consortium (ftp://ftp.1000genomes.ebi.ac.uk/vol1/ftp/data_collections/hgsv_sv_discovery/working/integration/20170515_Integrated_indels_Illumina_PacBio/Illumina_Indels_Merged_20170515.vcf.gz, ftp://ftp.1000genomes.ebi.ac.uk/vol1/ftp/data_collections/hgsv_sv_discovery/working/20180627_PhasedSVMSPAC/PhasedSVMsPAC.NA19240.vcf). Additionally, BioNano SV calls (ftp://ftp.1000genomes.ebi.ac.uk/vol1/ftp/data_collections/hgsv_sv_discovery/working/20180502_bionano/GM19240_DLE1_SV_hg38_indel.vcf and ftp://ftp.1000genomes.ebi.ac.uk/vol1/ftp/data_collections/hgsv_sv_discovery/working/20170109_BioNano_SV_update/GM19240_YRI_Daughter_20170109_bionano_SVMerged_InDel.vcf.gz) were used to validate SVs identified by IGenotyper in NA19240.

### Analysis of SNVs in regions accessible to next generation sequencing methods

SNVs were evaluated to determine if they were within regions accessible by next generation sequencing methods. The “strict” accessibility tracks were converted to bed format from: https://hgdownload.soe.ucsc.edu/gbdb/hg19/1000Genomes/phase3/20141020.strict_mask.whole_genome.bb.

The number of SNVs within the “strict” accessible regions was determined by using the bedtools “intersect” command.

### Calculating Hardy-Weinberg Equilibrium on NA19240 and NA12878 SNVs

SNVs from the 1000 Genomes Phase 3 project were downloaded from ftp://ftp.1000genomes.ebi.ac.uk/vol1/ftp/release/20130502/ALL.chr14.phase3_shapeit2_mvncall_integrated_v5a.20130502.genotypes.vcf.gz. Samples corresponding to the “EUR” superpopulation were extracted from the VCF file, and samples corresponding to “AFR” superpopulation were extracted into a separate VCF file. HWE was calculated using vcftools with the option “--hardy”.

## Supplementary Materials

### Methods

Fig. S1. Read length and average base call accuracy across CCS reads in CHM1.

Fig. S2. Three large discrepancies in CHM1 and NA19240 IGenotyper assemblies are expansions of 59mer tandem repeat motif.

Fig. S3. Parent-child trios used in study.

Fig. S4. V(D)J recombination in NA19240

Fig. S5. V(D)J recombination in NA12878

Fig. S6. Assembling all reads in regions of homozygosity

Fig. S7. Validation of insertion with IGHV7-4-1 gene in NA19240

Fig. S8. Validation of complex structural variant with IGHV1-8 and IGHV3-9 genes in NA19240

Fig. S9. Validation of duplication with IGHV3-23D gene in NA19240

Fig. S10. Validation of previously detected duplications harboring IGHV4-28, IGHV3-30, IGHV4-30-2, IGHV3-30-3, IGHV4-30-5, IGHV3-30-5, IGHV4-31, IGHV3-33 and IGHV4-34 genes

Fig. S11. Validation of insertion harboring IGHV4-38-23, IGHV3-43D, IGHV3-38-3 and IGHV1-38-4 gene.

Fig. S12. Validation of previously detected insertion harboring IGHV1-69, IGHV2-70D, IGHV1-69-2, IGH1-69D and IGHV2-70 genes.

Fig. S13. Analysis of SNVs in IGH that fail or pass HWE

Table S1. Sequences used to make custom IGH reference.

Table S2. Regions with designed capture probes

Table S3. Boosted probe regions

Table S4. Samples used in this study sequenced with different platforms and panels.

Table S5. Number and total bases of errors from incorrectly inserted sequence and missing sequence (indel errors) in the assembly of CHM1, NA19240 and NA12878.

Table S6. Indel errors in the assemblies separated by size with homopolymer annotation.

Table S7. Coordinates of V(D)J recombination in the two trios whose genomic DNA were derived from LCLs.

Table S8. Number of haplotype blocks and heterozygous blocks

Table S9. Number of fosmids used to validate assemblies.

Table S10. Mendelian inconsistencies rate in homozygous blocks Table S11. Embedded structural variants in the IG-reference.

Table S12. Alleles for IGHV genes for NA19240 and the inherited alleles in the parents of NA19240 (NA19238 and NA19239)

Table S13. Sequence for novel alleles detected in NA19240

Table S14. Alleles for IGHV genes for NA12878 and the inherited alleles in the parents of NA12878 (NA12892 and NA12891)

Table S15. Sequence for novel alleles detected in NA12878

Table S16. Validation of genotyped structural variants with fosmids and parental assemblies.

Table S17. Number of SNVs lifted over to GRCh37/hg19 in NA19240 and NA12878

Table S18. Number of overlapping SNVs in NA19240 and NA12878 between IGenotyper and the 1KGP phase 3 datasets

Table S19. Number of SNVs within accessible regions defined by 1KGP

Table S20. Number of SNVs passing or failing (p < 0.001) Hardy-Weinberg equilibrium

Table S21. Genotypes for NA19240 embedded structural variants in custom IGH reference. Table S22. Statistics from multiplexing replicates of NA12878

## Acknowledgments

This study makes use of a sample from an individual recruited by the Pacific Islands Rheumatic Heart Disease Genetics Network. This work was supported in part through the computational resources and staff expertise provided by the Scientific Computing at the Icahn School of Medicine at Mount Sinai.

## Author contributions

O.L.R., W.S.G., A.J.S., M.L.S., A.B. and C.T.W. conceived and planned the study. Computational analyses were performed by O.L.R. and T.P. Manuscript was written by O.L.R. and C.T.W. with contributions from T.P., E.E.E., A.J.S., W.S.G., M.L.S. and A.B. Code was written by O.L.R. Additional samples were provided by E.E.E., K.A. and T.P. Sequencing libraries were prepared by W.S.G., M.E. and M.L.S. Sequencing was performed by G.D, J.P., M.L.S. and R.S. C.T.W., A.B., and M.L.S. supervised the experiments, analysis, and data interpretation. All authors read and approved of the final paper.

