## Supplementary Text for "A novel framework for characterizing genomic haplotype diversity in the human immunoglobulin heavy chain locus"


Methods

Fig. S1. Read length and average base call accuracy across CCS reads in CHM1.

Fig. S2. Three large discrepancies in CHM1 and NA19240 IGenotyper assemblies are expansions of 59mer tandem repeat motif.

Fig. S3. Parent-child trios used in study.

Fig. S4. V(D)J recombination in NA19240

Fig. S5. V(D)J recombination in NA12878

Fig. S6. Assembling all reads in regions of homozygosity

Fig. S7. Validation of insertion with IGHV7-4-1 gene in NA19240

Fig. S8. Validation of complex structural variant with IGHV1-8 and IGHV3-9 genes in NA19240

Fig. S9. Validation of duplication with IGHV3-23D gene in NA19240

Fig. S10. Validation of previously detected duplications harboring IGHV4-28, IGHV3-30, IGHV4-30-2, IGHV3-30-3, IGHV4-30-5, IGHV3-30-5, IGHV4-31, IGHV3-33 and IGHV4-34 genes

Fig. S11. Validation of insertion harboring IGHV4-38-23, IGHV3-43D, IGHV3-38-3 and IGHV1-38-4 gene.

Fig. S12. Validation of previously detected insertion harboring IGHV1-69, IGHV2-70D, IGHV1-69-2, IGH1-69D and IGHV2-70 genes.

Fig. S13. Analysis of SNVs in IGH that fail or pass HWE

Table S1. Sequences used to make custom IGH reference.

Table S4. Samples used in this study sequenced with different platforms and panels.

Table S5. Number and total bases of errors from incorrectly inserted sequence and missing sequence (indel errors) in the assembly of CHM1, NA19240 and NA12878.

Table S6. Indel errors in the assemblies separated by size with homopolymer annotation.

Table S7. Coordinates of V(D)J recombination in the two trios whose genomic DNA were derived from LCLs.

Table S8. Number of haplotype blocks and heterozygous blocks

Table S9. Number of fosmids used to validate assemblies.

Table S10. Mendelian inconsistencies rate in homozygous blocks

Table S11. Embedded structural variants in the IG-reference.

Table S12. Alleles for IGHV genes for NA19240 and the inherited alleles in the parents of NA19240 (NA19238 and NA19239)

Table S13. Sequence for novel alleles detected in NA19240

Table S14. Alleles for IGHV genes for NA12878 and the inherited alleles in the parents of NA12878 (NA12892 and NA12891)

Table S15. Sequence for novel alleles detected in NA12878

Table S16. Validation of genotyped structural variants  with fosmids and parental assemblies.

Table S17. Number of SNVs lifted over to GRCh37/hg19 in NA19240 and NA12878

Table S18. Number of overlapping SNVs in NA19240 and NA12878 between IGenotyper and the 1KGP phase 3 datasets

Table S19. Number of SNVs within accessible regions defined by 1KGP

Table S20. Number of SNVs passing or failing (p < 0.001) Hardy-Weinberg equilibrium

Table S21. Genotypes for NA19240 embedded structural variants in custom IGH reference.

Table S22. Statistics from multiplexing replicates of NA12878

### Methods

#### Description of capture panels

Three different target panels ("A", "B" and "C") were developed and tested. The sequencing probes were developed by providing Roche with a fasta file representing IGH sequence targets. Roche then returned coordinates of targeted regions. Targets for panel A were developed from 12 different fosmid sequences spanning the IGH locus and regions within chromosome 14 of hg38 (Table S1). Targets for panel B were developed from the custom IGH reference we created for IGenotyper (Figure 1a). Targets for panel C were developed from the custom IGH reference and 97 additional regions from hg38. Ninety-six additional regions correspond to Ancestry Informative Markers (AIMs). The last additional region corresponds to the IGHC region. Analysis features for AIMs and the IGHC region are currently not operational in IGenotyper, but will be incorporated into future versions.

In panel B and C, additional probes (boosting) were added to increase the sequencing coverage (Table S3). The amount of boosting ranged from 2 to 5. The mean coverage was consistent in panel B across IGH, we noted a loss in coverage over the IGHJ region in panel A. We speculate this is caused by a lack of adjacent target sequence on the 3' flank of the IGHJ region in panel A, in contrast to panel B, which also included sequence targets across the entirety of the IGHC region.For CHM1, we combined data from panels A and B to mitigate inconsistencies in regional coverage between them.

Sequence target files will be made available by request.

**Large insertions within the IGenotyper assemblies not found in GRCh38 and fosmids**

Haplotype 1 of NA19240 contained a 1,704 bp and 1,456 bp insertion that was not found in the sequenced fosmids. The CHM1 assembly contained a 2,226 bp insertion not found in GRCh38. These were the largest differences that were found between the IGenotyper assemblies and the ground truth sequences. All three large insertions were found within the duplication containing the genes *IGHV1-69, IGHV2-70, IGHV1-69-2, IGHV1-69D* and *IGHV2-70D*. All these insertions contained an expansion of the tandem repeat motif "TTTAAAAAGATAGTTCTCATCGCCTTGAATTGTGGGAGCAGCTCAGATGTGATAGAATA". This tandem repeat has been previously difficult to assemble. The BAC clone CH17-212P11 (AC245369.4) underlying this region in GRCh38 did not resolve this duplication. Under features in the genbank accession, the tandem repeat is labelled as "unresolved duplication". Therefore it is possible that the IGenotyper assembly is correct, and the reference is incorrect.

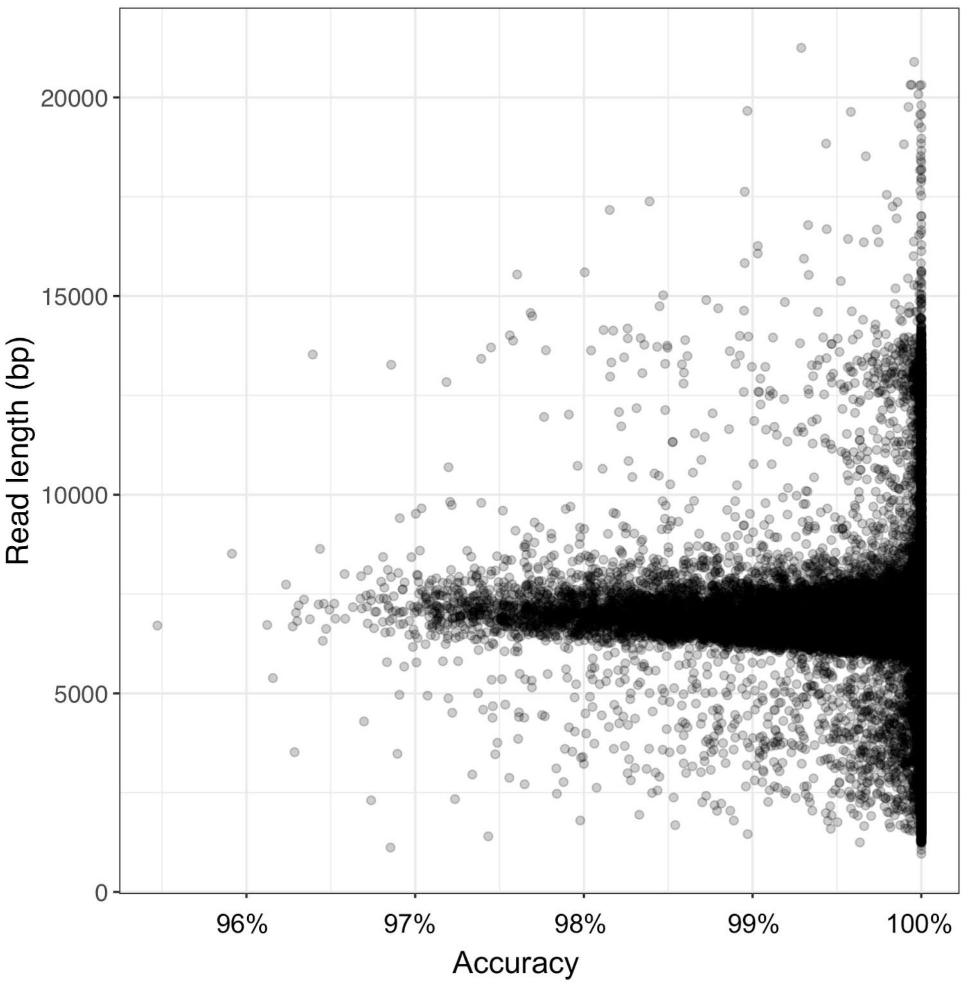


**Fig. S1. Read length and average base call accuracy across CCS reads in CHM1.**

Every point is a CCS read. Raw SMRT sequences with at least two passes were converted into CCS reads. The phred quality score for every base in the CCS read was averaged. The y-axis has the base call accuracy of the average phred quality score. The x-axis is the length of the CCS read.

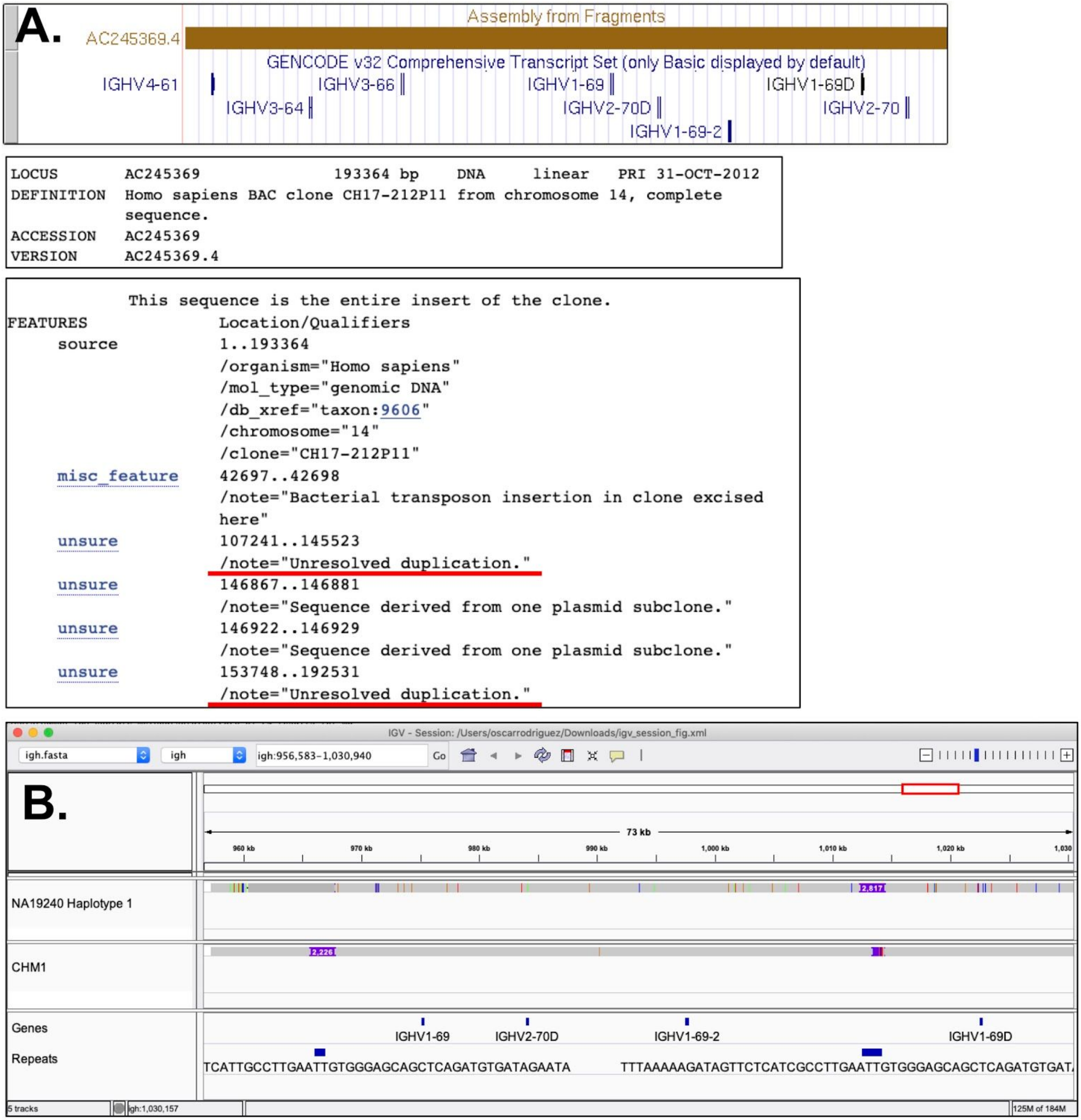


**Fig. S2. Three large discrepancies in CHM1 and NA19240 IGenotyper assemblies are expansions of 59mer tandem repeat motif.**

(**A**) The BAC clone, AC245369.4, was used to assemble the IGH region containing the genes from *IGHV3-66* to *IGHV2-70*. The BAC clone contained two unresolved duplications (red lines). The unresolved duplications are a tandem repeat with a 56-mer motif

"TTTAAAAAGATAGTTCTCATCGCCTTGAATTGTGGGAGCAGCTCAGATGTGATAGAATA" shown in (**B**).

(**B**) The CHM1 assembly from IGenotyper contained two large insertion sequences of 1,704 bp and 1,456 bp within this tandem repeat. Since the duplication was not resolved in GRCh38, we are uncertain if IGenotyper is correct or not. The NA19240 haplotype 1 also contains an insertion with this tandem repeat in the IGenotyper assembly that does not agree with the NA19240 haplotype 1 fosmids spanning this region. This can be due to misassembly, somatic expansion, or an artifact in the cell line.

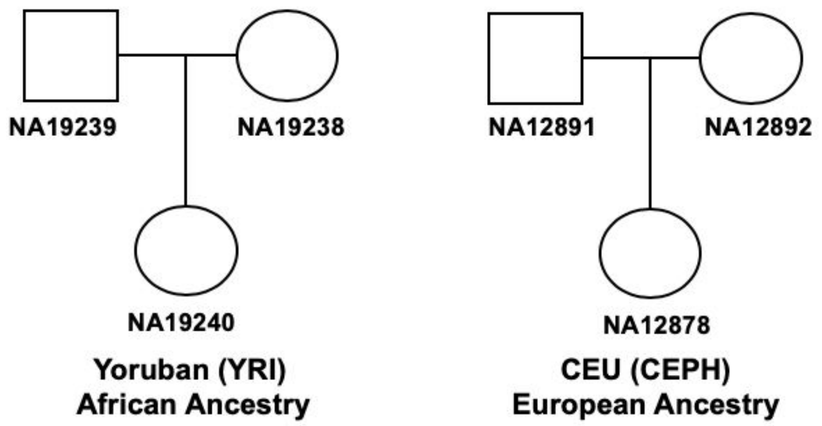


**Fig. S3. Parent-child trios used in study.**

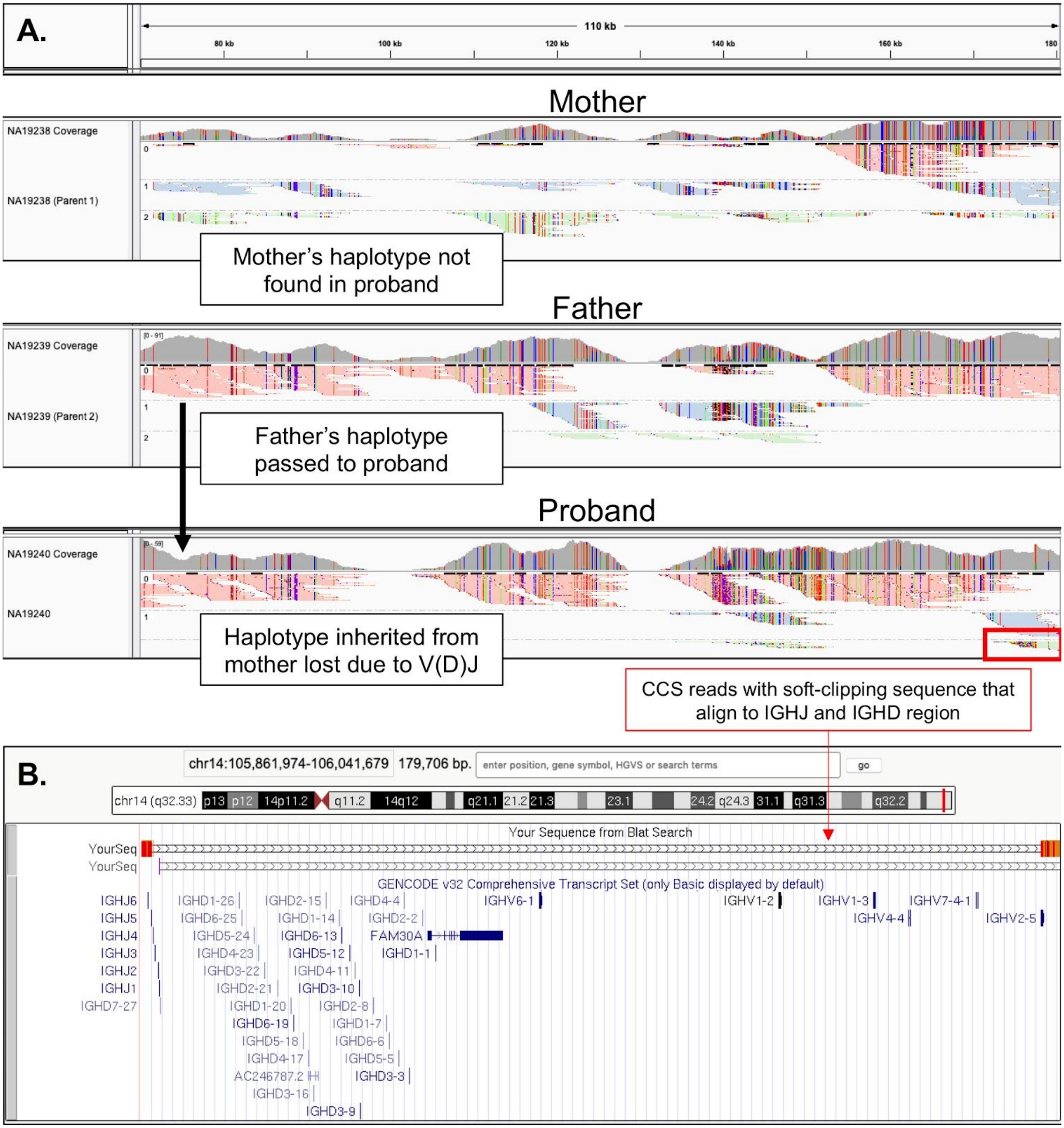


**Fig. S4. V(D)J recombination in NA19240**

**(A)** IGV screenshot of CCS reads from NA19240, NA19238 and NA19239 aligned to the proximal region of IGH. NA19240 (proband) lost the beginning part of a single haplotype due to V(D)J recombination. The proband still contains the haplotype inherited from the father (NA19239) but not the mother's inherited haplotype (NA19238). The mother's inherited haplotype appears after *IGHV2-5* in the proband. CCS reads spanning *IGHV2-5* in the proband are not fully aligned to the reference and contain soft-clipping sequences that align to the D and J genes as shown in the BLAT output in (**B**).

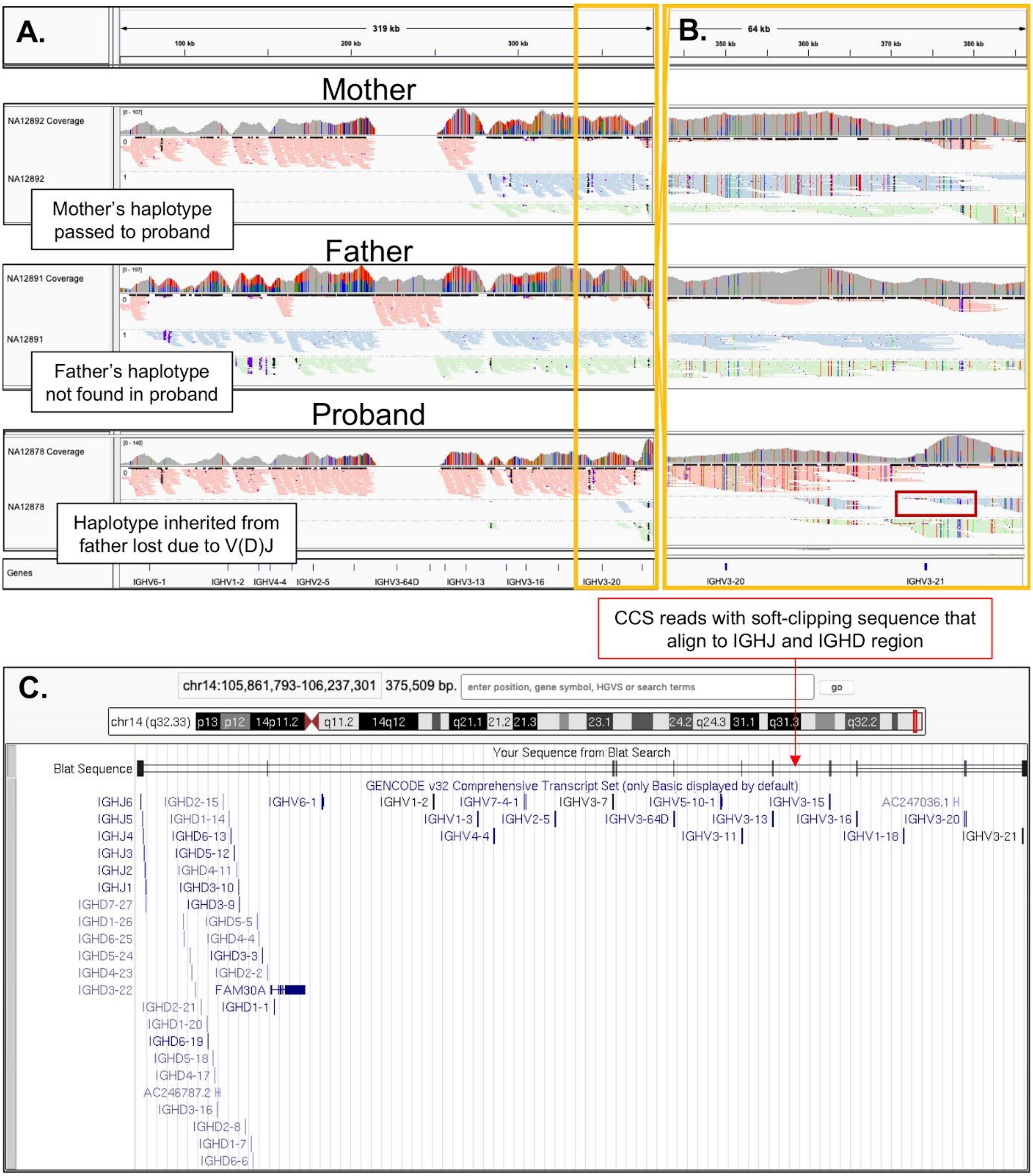


**Fig. S5. V(D)J recombination in NA12878**

**(A,B)** IGV screenshot of CCS reads from NA12878, NA12891 and NA12892 aligned to the proximal region of IGH. NA12878 (proband) lost the beginning part of a single haplotype due to V(D)J recombination. The proband still contains the haplotype inherited from the mother (NA12892) but not the father's inherited haplotype (NA12891). The father's inherited haplotype appears after *IGHV3-21* in the proband. CCS reads spanning *IGHV3-21* in the proband are not fully aligned to the reference and contain soft-clipping sequences that align to the D and J genes as shown in the BLAT output in (**B**).

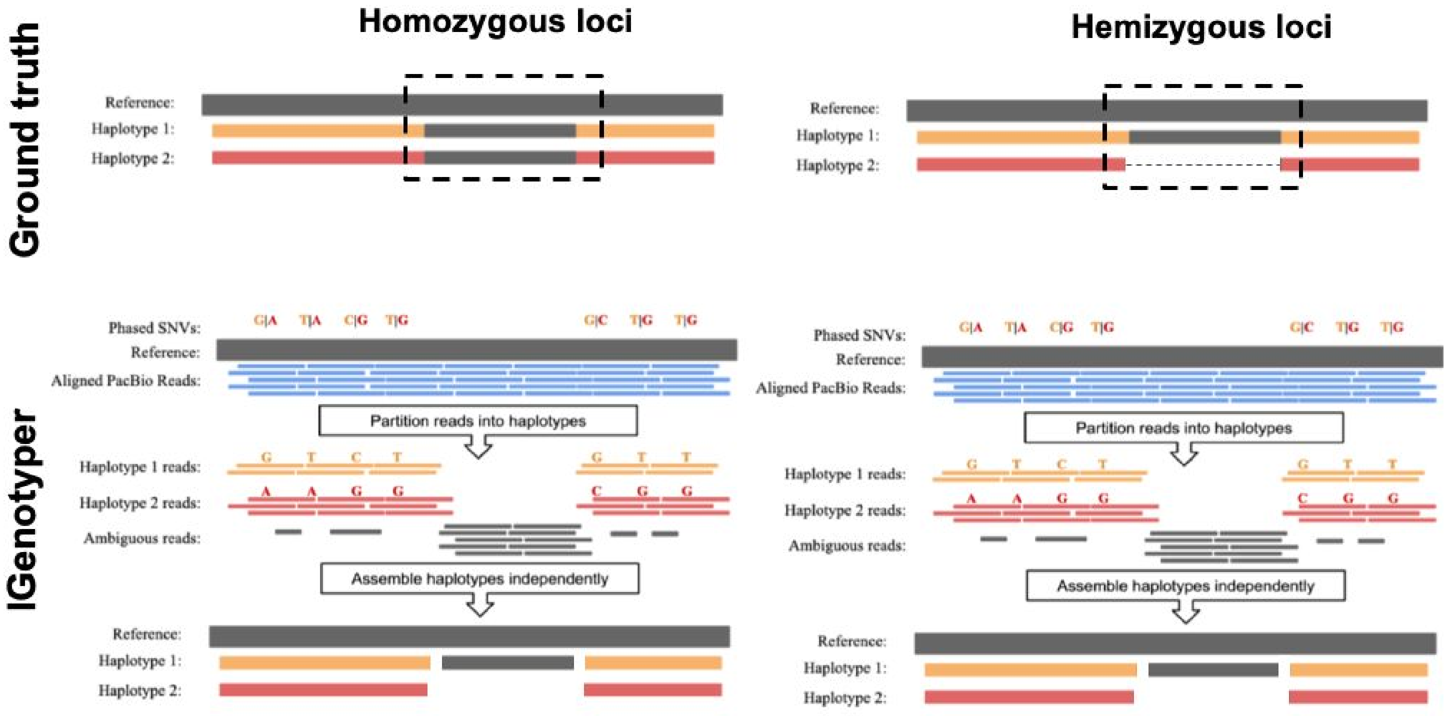


**Fig. S6. Assembling all reads in regions of homozygosity**

Schematic showing the phased reads and assembly process in a region of homozygosity and hemizygosity. All reads in both cases are assembled together. The assembly process is the same.

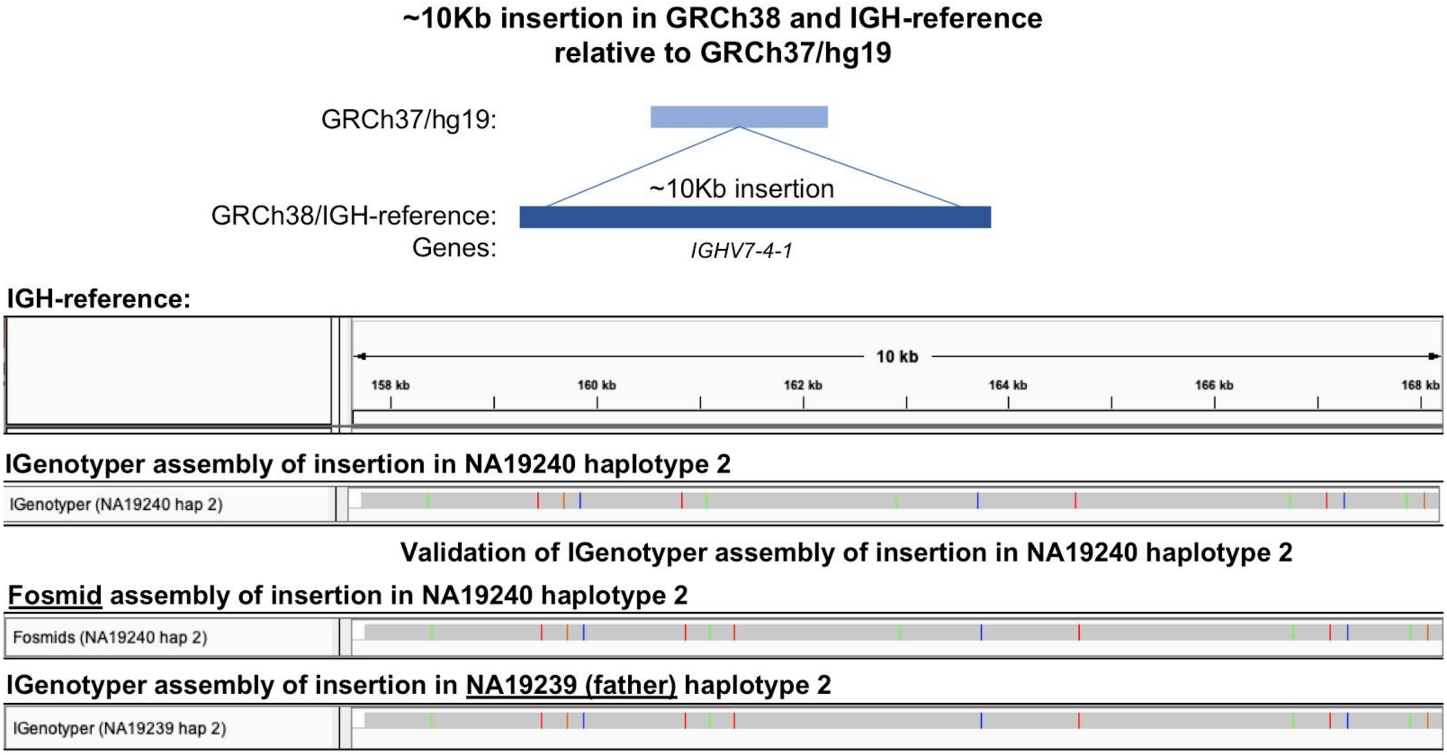


**Fig. S7. Validation of insertion with *IGHV7-4-1* gene in NA19240**

Top: Schematic showing insertion identified by Watson et al 2013 in CHM1 (GRCh38) relative to GRCh37/hg19. The insertion from CHM1 is present in the IGH-reference.

Bottom: IGenotyper assemblies and fosmids shown in order:

1) NA19240 haplotype 2
2) NA19240 haplotype 2 fosmids
3) NA19239 haplotype 2

NA19240 contained the insertion in a single haplotype. The alternate haplotype was lost due to a V(D)J event. The insertion was validated through fosmids and inheritance with parental data. The assembly of the insertion shows high concordance with sequenced fosmids and the assembled haplotype from NA19239 (father).

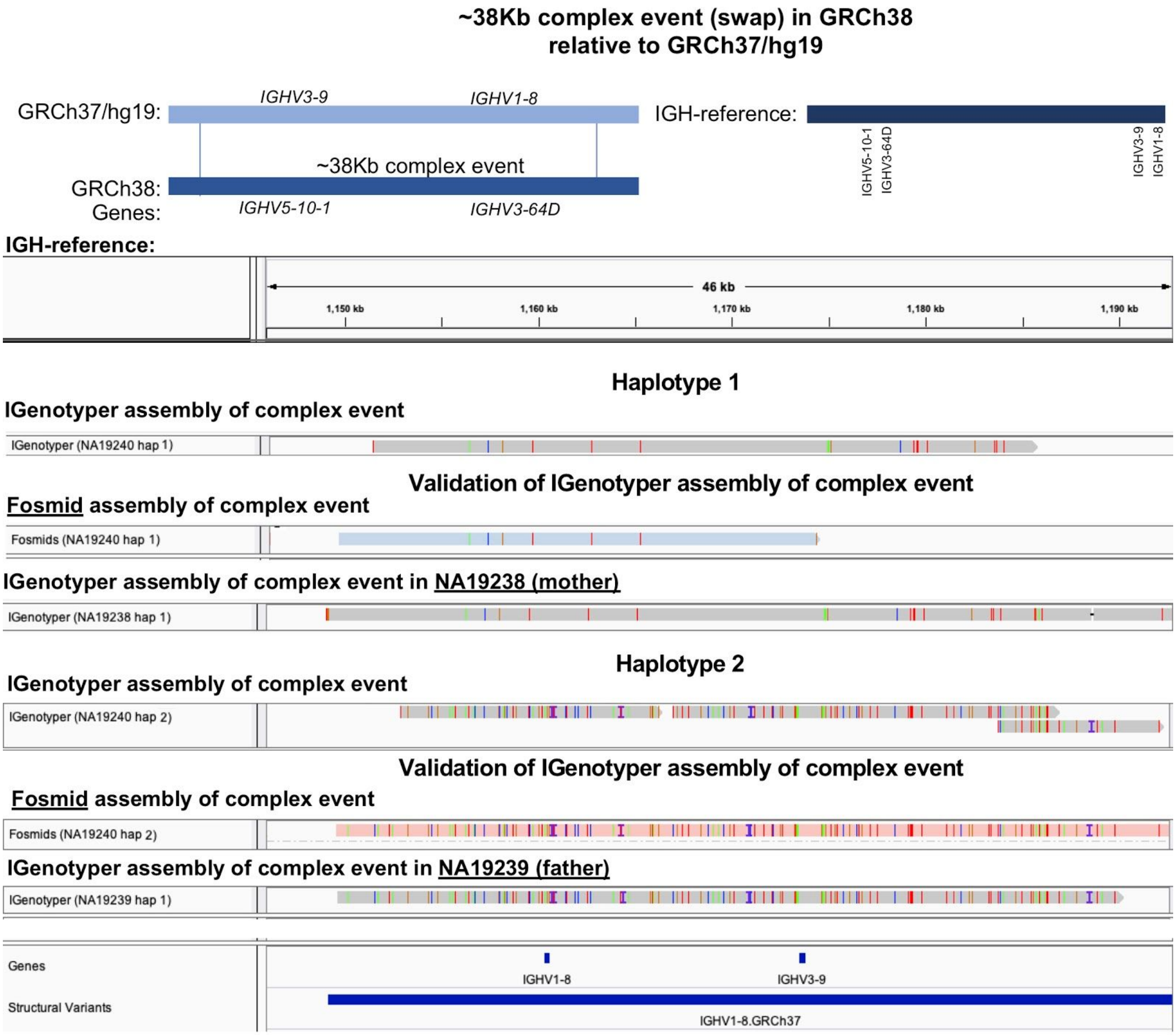


**Fig. S8. Validation of complex structural variant with *IGHV1-8* and *IGHV3-9* genes in NA19240**

Top: Schematic showing the complex event identified by Watson et al 2013 in CHM1 (GRCh38) relative to GRCh37/hg19. The complex event is not a straightforward insertion or deletion, rather it is a swap of sequence of the same length. The sequence present in GRCh37/hg19 was placed at the end of the IGH-reference (right).

Bottom: IGenotyper assemblies and fosmids shown in order:

1) NA19240 haplotype 1
2) NA19240 haplotype 1 fosmids
3) NA19238 haplotype 1

4) NA19240 haplotype 2
5) NA19240 haplotype 2 fosmids
6) NA19239 haplotype 1

NA19240 contained the GRCh37/hg19 haplotype in both its haplotype (1 and 2). Both haplotypes were validated by parental data. Fosmid sequence validated both haplotypes and parental data partially validated haplotype 1 and fully validated haplotype 2.

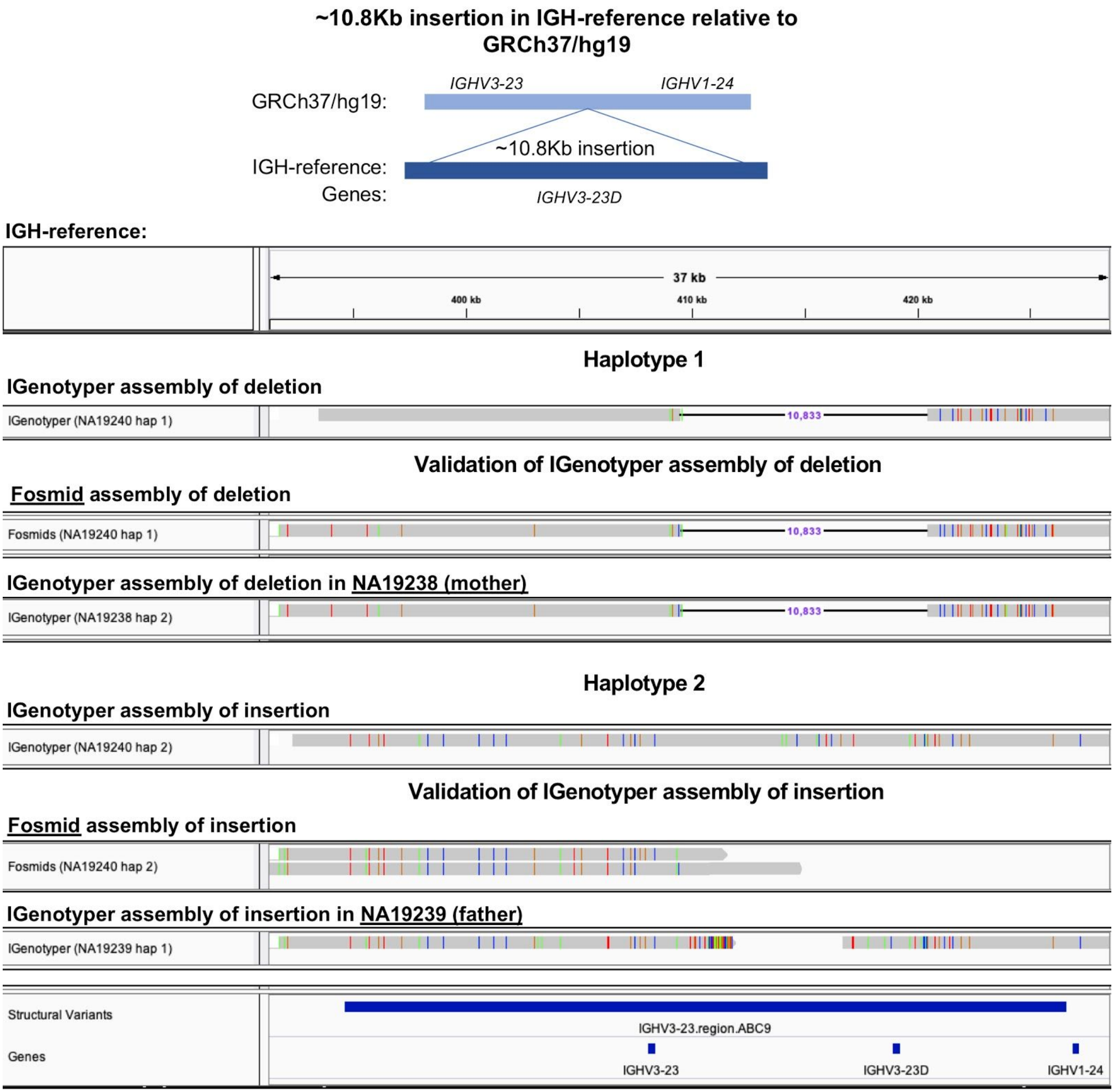


**Fig. S9. Validation of duplication with *IGHV3-23D* gene in NA19240**

<u>Top</u>: Schematic showing the duplication identified by Watson et al 2013 in NA18956 and NA12156 relative to GRCh37/hg19. The duplication was inserted in the IGH-reference.

<u>Bottom</u>: IGenotyper assemblies and fosmids shown in order:

1) NA19240 haplotype 1
2) NA19240 haplotype 1 fosmids

3) NA19238 haplotype 2

4) NA19240 haplotype 2

5) NA19240 haplotype 2 fosmids

6) NA19239 haplotype 1

NA19240 contained the insertion in one haplotype (2). The absence of the duplication in the alternate haplotype was validated by fosmids and parental data. The presence of the duplication was partially validated by fosmids and parental data.

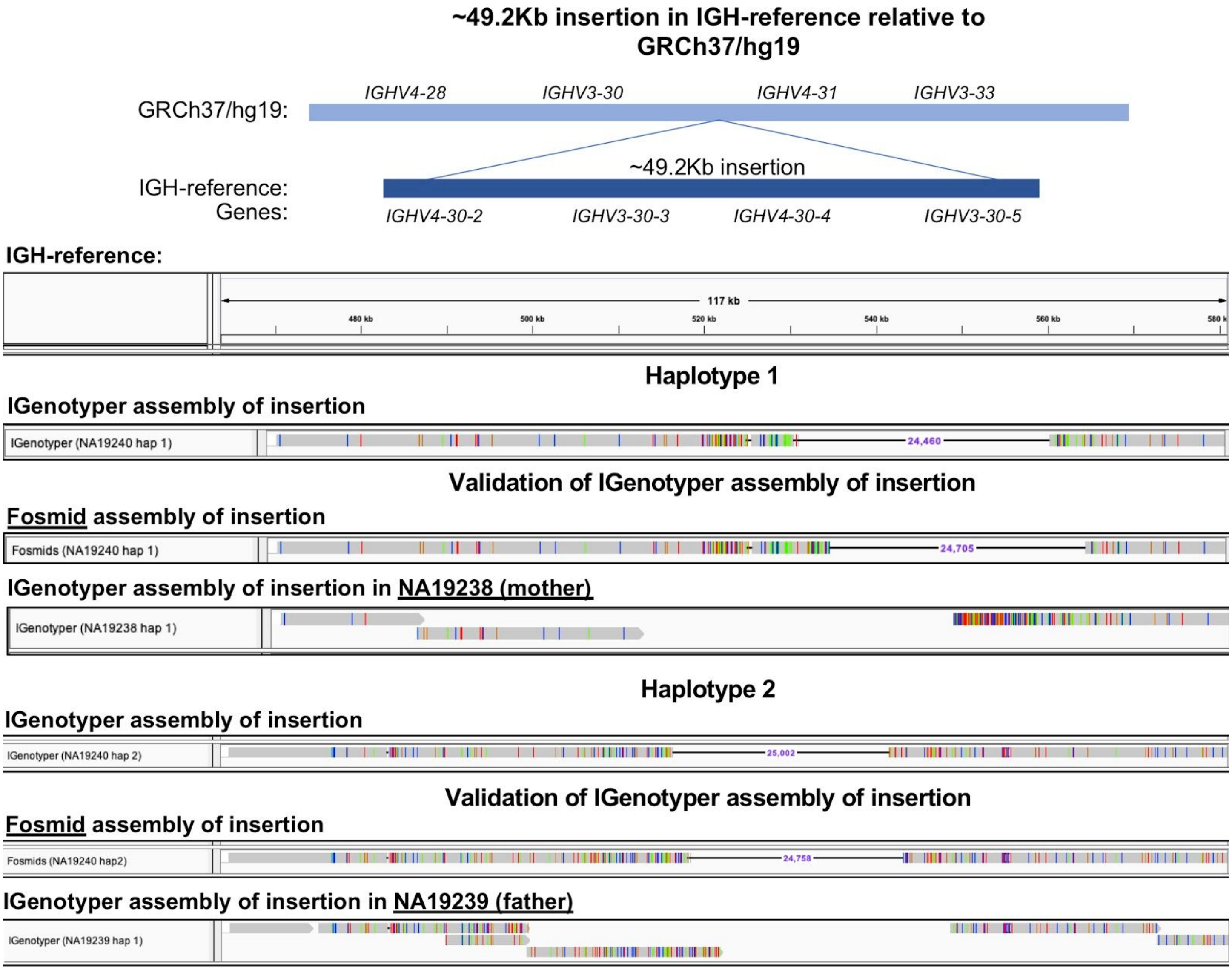

**Fig. S10. Validation of previously detected duplications harboring *IGHV4-28, IGHV3-30, IGHV4-30-2, IGHV3-30-3, IGHV4-30-5, IGHV3-30-5, IGHV4-31, IGHV3-33* and *IGHV4-34* genes**

Top: Schematic showing the duplication identified by Watson et al 2013 in NA18555 relative to GRCh37/hg19. The duplication was inserted in the IGH-reference.

Bottom: IGenotyper assemblies and fosmids shown in order:

1) NA19240 haplotype 1
2) NA19240 haplotype 1 fosmids
3) NA19238 haplotype 2
4) NA19240 haplotype 2
5) NA19240 haplotype 2 fosmids
6) NA19239 haplotype 1

NA19240 contained a partial insertion in haplotype 1 and 2. The partial insertions were validated by fosmids and partially by parental data. The IGenotyper assembly of haplotype 1 (1) and the fosmids of haplotype 1 (2) have a high sequence concordance (>99.99%) but due to small gaps and small number of base mismatches BLASR (the mapping aligner) shift the ~24 Kbp deletion (Table S16).

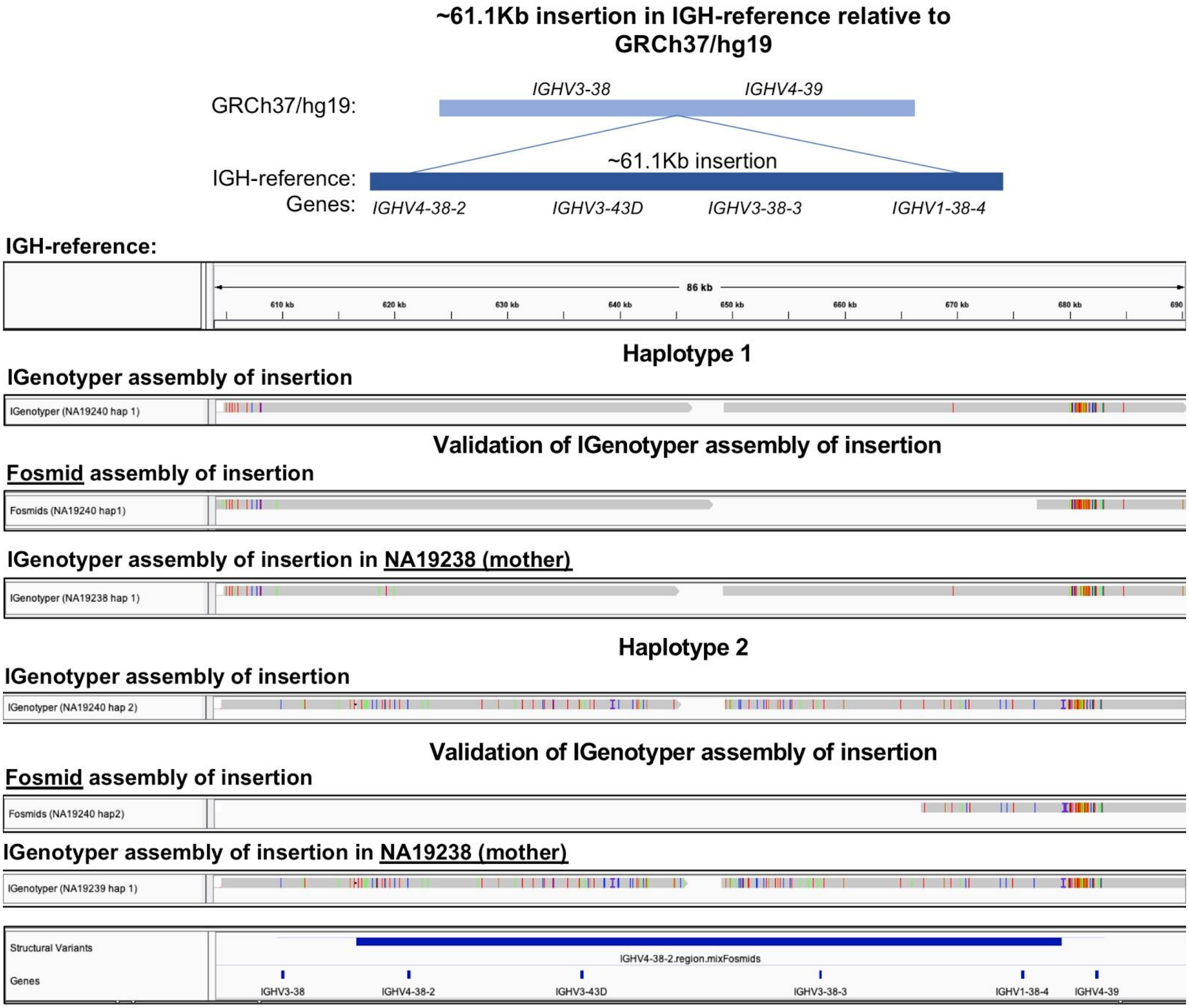


**Fig. S11. Validation of insertion harboring *IGHV4-38-23, IGHV3-43D, IGHV3-38-3* and *IGHV1-38-4* gene.**

Top: Schematic showing the duplication identified by Watson et al 2013 in NA15510 and NA19240 relative to GRCh37/hg19. The duplication was inserted in the IGH-reference.

Bottom: IGenotyper assemblies and fosmids shown in order:

1) NA19240 haplotype 1
2) NA19240 haplotype 1 fosmids
3) NA19238 haplotype 2

4) NA19240 haplotype 2

5) NA19240 haplotype 2 fosmids

6) NA19239 haplotype 1

NA19240 contained the insertion in both haplotypes. Both insertions were validated with parental data and partially validated by fosmids.

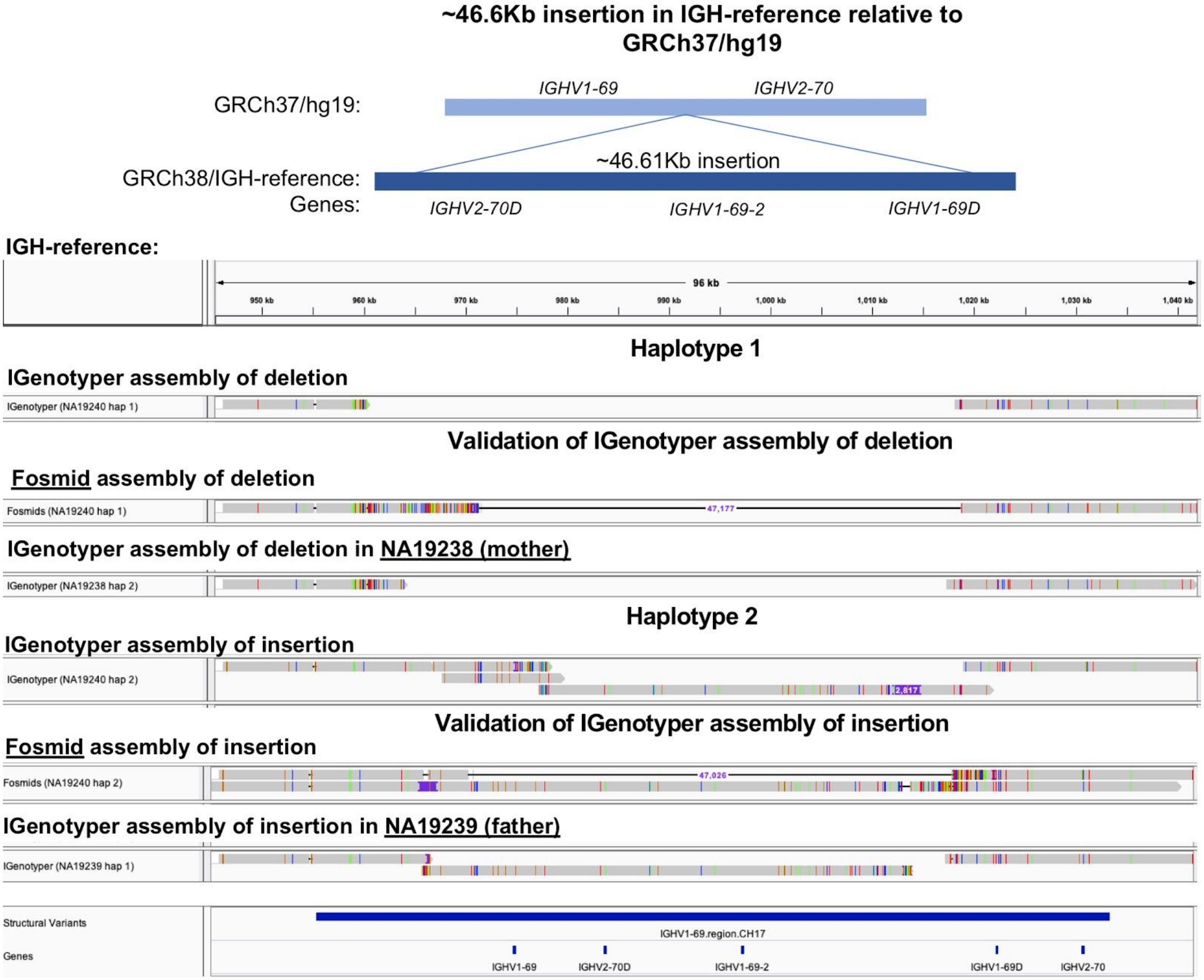

**Fig. S12. Validation of previously detected insertion harboring *IGHV1-69, IGHV2-70D, IGHV1-69-2, IGH1-69D* and *IGHV2-70* genes.**

<u>Top</u>: Schematic showing the insertion identified by Watson et al 2013 in CHM1 (GRCh38) relative to GRCh37/hg19. The insertion is present in the IGH-reference.

<u>Bottom</u>: IGenotyper assemblies and fosmids shown in order:

1) NA19240 haplotype 1
2) NA19240 haplotype 1 fosmids
3) NA19238 haplotype 2
4) NA19240 haplotype 2
5) NA19240 haplotype 2 fosmids
6) NA19239 haplotype 1

The insertion was present only in a single haplotype (2). The absence of the insertion was validated by fosmids and parental data. The insertion was validated by fosmids and parental data. There is evidence both in the CCS from the target-enrichment/IGenotyper sequencing run and fosmid clones that the insertion haplotype underwent a somatic mutation. The insertion haplotype passed on by the father (NA19239) was assembled as an insertion and deletion with the fosmids.

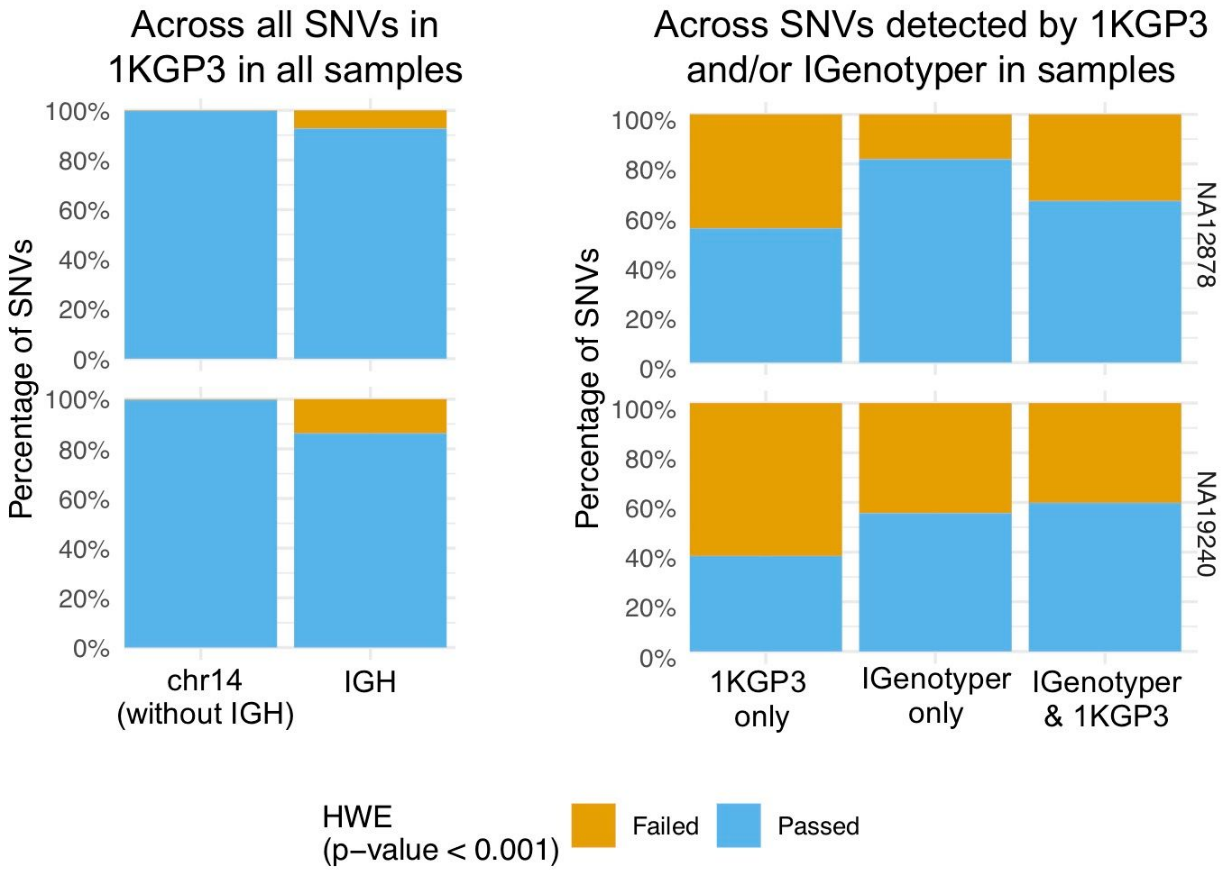


**Fig. S13. Analysis of SNVs in IGH that fail or pass HWE**

As a reference point, the percentage of SNVs in the 1KGP3 dataset for European samples (top left) and African samples (bottom left) in chromosome 14, excluding SNVs in the IGH locus, that fail or pass HWE was calculated. The percentage of SNVs in IGH for the same samples across all the SNVs in the 1KGP3 dataset that failed or passed HWE was also calculated. The SNVs were subsetted to those that:

1. Were found only in the 1KGP3 dataset for NA12878 and NA19240
2. Were found only by IGenotyper
3. Were found both by IGenotyper and in the 1KGP3

The percentage of SNVs that failed or passed HWE for each subset was calculated across all the African or European samples in the 1KGP3 dataset.

**Table S1. Sequences used to make custom IGH reference.**

| **IGH reference** | | | **Source sequence** | | | |
|---|---|---|---|---|---|---|
| **chrom** | **start** | **end** | **chrom** | **start** | **end** | **source** |

| | | | | | | |
|---|---|---|---|---|---|---|
| igh | 1 | 394,082 | chr14 | 105,860,500 | 106,254,581 | GRCh38 |
| igh | 394,082 | 412,363 | AC206018.3 | 21,438 | 39,718 | ABC9-43993300H10 |
| igh | 412,363 | 427,266 | AC244473.3 | 2,961 | 17,864 | ABC9-43849600N9 |
| igh | 427,266 | 484,858 | chr14 | 106,276,923 | 106,317,171 | GRCh38 |
| igh | 484,858 | 519,568 | AC231260.2 | 3,653 | 38,382 | ABC11-47150400I4 |
| igh | 519,568 | 548,278 | AC244456.2 | 11,919 | 40,628 | ABC11-47354200D2 |
| igh | 548,278 | 562,484 | KC162925.1 | 19,397 | 33,602 | ABC11-49598600E10 |
| igh | 562,484 | 609,357 | chr14 | 106,363,211 | 106,403,456 | GRCh38 |
| igh | 609,357 | 632,150 | KC162926.1 | 19,561 | 42,354 | ABC10-44084700I10 |
| igh | 632,150 | 662,842 | AC233755.2 | 13,040 | 43,732 | ABC10-44145400L1 |
| igh | 662,842 | 685,205 | AC241995.3 | 7,463 | 29,824 | WI2-1707G1 |
| igh | 685,206 | 1,144,129 | chr14 | 106,424,795 | 106,883,718 | GRCh38 |
| igh | 1,144,130 | 1,149,130 | NA | NA | NA | Gap sequence: "N" |
| igh | 1,149,131 | 1,193,129 | chr14 | 106,527,905 | 106,571,904 | hg19 |

**Table S4. Samples used in this study sequenced with different platforms and panels.**

| **Sample** | **Replicate** | **Genomic DNA Source** | **Capture design** | **Approximate Insert size (kb)** | **PacBio Platform** | **# of SMRT cells** | **Multiplexed** | **Population/ Ethnicity Information** |
|---|---|---|---|---|---|---|---|---|
| CHM1 | No | Hydatidiform Mole Cell Line | A | 6 | RSII | 2 | No | Caucasian |
| CHM1 | No | Hydatidiform Mole Cell Line | B | 6 | Sequel | 1 | No | Caucasian |
| CHM1 | No | Hydatidiform Mole Cell Line | A+B | 6 | . | . | . | Caucasian |
| NA19238 | No | Lymphoblastoid Cell Line | C | 7.5 | Sequel 1 | 1 | Yes (5 samples) | YRI |
| NA19239 | No | Lymphoblastoid Cell Line | C | 7.5 | Sequel 1 | 1 | Yes (5 samples) | YRI |
| NA19240 | No | Lymphoblastoid Cell Line | B | 7.5 | RSII | 2 | No | YRI |
| NA12891 | No | Lymphoblastoid Cell Line | C | 7.5 | Sequel 1 | 1 | Yes (5 samples) | CEU |

| | | | | | | | | |
|---|---|---|---|---|---|---|---|---|
| NA12892 | No | Lymphoblastoid Cell Line | C | 7.5 | Sequel 1 | 1 | Yes (5 samples) | CEU |
| NA12878 | No | Lymphoblastoid Cell Line | C | 7.5 | Sequel 1 | 1 | Yes (5 samples) | CEU |
| NA12878 | Yes | Lymphoblastoid Cell Line | B | 7.5 | Sequel 1 | 1 | Yes (8 samples) | CEU |
| NA12878 | Yes | Lymphoblastoid Cell Line | B | 7.5 | Sequel 1 | 1 | Yes (8 samples) | CEU |
| NA12878 | Yes | Lymphoblastoid Cell Line | B | 7.5 | Sequel 1 | 1 | Yes (8 samples) | CEU |
| NA12878 | Yes | Lymphoblastoid Cell Line | B | 7.5 | Sequel 1 | 1 | Yes (8 samples) | CEU |
| NA12878 | Yes | Lymphoblastoid Cell Line | B | 7.5 | Sequel 1 | 1 | Yes (8 samples) | CEU |
| NA12878 | Yes | Lymphoblastoid Cell Line | B | 7.5 | Sequel 1 | 1 | Yes (8 samples) | CEU |
| NA12878 | Yes | Lymphoblastoid Cell Line | B | 7.5 | Sequel 1 | 1 | Yes (8 samples) | CEU |
| NA12878 | Yes | Lymphoblastoid Cell Line | B | 7.5 | Sequel 1 | 1 | Yes (8 samples) | CEU |
| Sample A (Parks et al) | No | Peripheral Blood Mononuclear Cells | A | 6 | RSII | 2 | No | Fijian |

**Table S5. Number and total bases of errors from incorrectly inserted sequence and missing sequence (indel errors) in the assembly of CHM1, NA19240 and NA12878.**

| Sample | Assembly size (bp) | Indel errors | Total missing sequence (bp) | Total inserted sequence (bp) | Percentage of missing sequence | Percentage of inserted sequence |
|---|---|---|---|---|---|---|
| CHM1 | 1,009,792 | 220 | 132 | 2,840 | 0.01% | 0.28% |
| NA19240 (diploid) | 1,829,616 | 276 | 217 | 3,984 | 0.01% | 0.22% |
| NA12878 (diploid) | 1,442,310 | 188 | 123 | 626 | 0.01% | 0.04% |

**Table S6. Indel errors in the assemblies separated by size with homopolymer annotation.**

| Sample | Gaps greater than 100bp | Gaps between 2bp and 100bp | 1bp or 2bp gaps |
|---|---|---|---|

| | Del | Del bases (bp) | Ins | Ins bases (bp) | Del | Del bases (bp) | Ins | Ins bases (bp) | Del | Del bases (bp) | Ins | Ins bases (bp) | Del/ins in homopolymers |
|---|---|---|---|---|---|---|---|---|---|---|---|---|---|
| CHM1 | 0 | 0 | 3 | 2,521 | 2 | 6 | 16 | 222 | 113 | 126 | 86 | 97 | 123 |
| NA19240 | 0 | 0 | 6 | 3,803 | 1 | 4 | 14 | 113 | 192 | 213 | 62 | 68 | 181 |
| NA12878 | 0 | 0 | 1 | 520 | 3 | 10 | 4 | 16 | 105 | 113 | 75 | 90 | 114 |

**Table S7. Coordinates of V(D)J recombination in the two trios whose genomic DNA were derived from LCLs.**

| **Sample** | **Chrom** | **Start** | **End** | **V(D)J gene** | **IGHV genes from single haplotype** | **Amount of IGHV region lost in one haplotype starting from IGHV6-1 (bp)** | **Amount of IGHV region from both haplotypes (bp)** | **Percentage of IGHV region from both haplotypes** |
|---|---|---|---|---|---|---|---|---|
| NA19240 | igh | 1 | 177,702 | *IGHV2-5* | *IGHV6-1, IGHV1-2, IGHV1-3, IGHV4-4, IGHV7-4-1* | 98,447 | 1,016,427 | 91.17% |
| NA19238 | NA | NA | NA | None | None | 0 | 1,114,874 | 100.00% |
| NA19239 | igh | 1 | 126,377 | *IGHV1-2* | *IGHV6-1, IGHV1-2* | 47,122 | 1,067,752 | 95.77% |
| NA12878 | igh | 1 | 374,857 | *IGHV3-21* | *IGHV6-1, IGHV1-2, IGHV1-3, IGHV4-4, IGHV7-4-1, IGHV2-5, IGHV3-11, IGHV3-13, IGHV3-15, IGHV3-16, IGHV1-18, IGHV3-20, IGHV1-8, IGHV3-9* | 295,602 | 819,272 | 73.49% |

| | | | | | | | | |
|---|---|---|---|---|---|---|---|---|
| NA12891 | igh | 1 | 79,558 | *IGHV6-1* | *IGHV6-1* | 303 | 1,114,571 | 99.97% |
| NA12892 | igh | 1 | 269,332 | *IGHV3-13* | *IGHV6-1, IGHV1-2, IGHV1-3, IGHV4-4, IGHV7-4-1, IGHV2-5, IGHV3-11, IGHV3-13* | 190,077 | 924,797 | 82.95% |

**Table S8. Number of haplotype blocks and heterozygous blocks**

| Sample | Haplotype blocks | # of heterozygous blocks | IGH reference bases in heterozygous blocks (bp) |
|---|---|---|---|
| NA19240 | 41 | 20 | 826,548 |
| NA12878 | 49 | 24 | 424,834 |

**Table S9. Number of fosmids used to validate assemblies.**

| Sample | Sequencing technology | Number of fosmids | Bases in assembly covered by fosmids | % of assembly covered by fosmid |
|---|---|---|---|---|
| NA19240 | PacBio | 85 | 1,505,709 | 82.30% |
| NA19240 | Sanger | 7 | 210,438 | 11.50% |
| NA12878 | PacBio | 73 | 1,180,299 | 81.83% |
| NA12878 | Sanger | 2 | 70,208 | 4.87% |

**Table S10. Mendelian inconsistencies rate in homozygous blocks**

| Sample | Mendelian inconsistencies | Bases from homozygous blocks | Mendelian inconsistencies rate |
|---|---|---|---|
| NA19240 | 27 | 57,313 | 0.047% |
| NA12878 | 23 | 139,029 | 0.017% |

**Table S11. Embedded structural variants in the IG-reference.**

| SV ID | chrom | start | end | Length | Embedded structural variant | IGHV genes |
|---|---|---|---|---|---|---|
| SV_1 | igh | 157,688 | 167,669 | 9,981 | Insertion | V7-4-1 |
| SV_2A* | igh | 210,158 | 257,471 | 47,313 | Complex event | V3-64D, V5-10-1 |
| SV_3 | igh | 394,658 | 426,627 | 31,969 | Duplication | V3-23, V3-23D |
| SV_4** | igh | 484,922 | 559,858 | 74,936 | Duplication | V4-28,V3-30,V4-30-2,V3-30-3,V4-30 |

| | | | | | | |
|---|---|---|---|---|---|---|
| | | | | | | -4,V3-30-5,V4-31,V3-33,V4-34 |
| SV_5 | igh | 609,413 | 682,906 | 73,493 | Insertion | V3-38,V4-38-2,V3-43D,V3-38-3,V4-39,V1-38-4 |
| SV_6 | igh | 955,770 | 1,033,787 | 78,017 | Insertion | V1-69,V2-70D,V1-69-2,V2-69D,V2-70 |
| SV_2B* | igh | 114,9129 | 1,194,129 | 45,000 | Complex event | V1-8,V3-9 |

*These complex events represent the same structural variant; on any given chromosome either V1-8/3-9 or V3-64/5-10-1 will be present.
**Given the complexity of this region, and known structural haplotype hypervariability, explicit SV genotype calls are not provided by IGenotyper for this SV.

**Table S12. Alleles for IGHV genes for NA19240 and the inherited alleles in the parents of NA19240 (NA19238 and NA19239)**

| Gene | Haplotype 1 | Haplotype 2 | NA19238 | NA19239 |
|---|---|---|---|---|
| IGHV6-1 | Allele lost by VDJ | *01 | Allele not needed | *01 |
| IGHV1-2 | Allele lost by VDJ | *02 | Allele not needed | *02 |
| IGHV1-3 | Allele lost by VDJ | *03[a] | Allele not needed | *03[a] |
| IGHV4-4 | Allele lost by VDJ | *08 | Allele not needed | *08 |
| IGHV7-4-1 | Allele lost by VDJ | *02 | Allele not needed | *02 |
| IGHV2-5 | Allele lost by VDJ | *02 | Allele not needed | *02 |
| IGHV3-7 | *01 | *01 | *01 | *01 |
| IGHV3-11 | *01 | *04 | *01 | *04 |
| IGHV3-13 | *01 | *03 | *01 | *03 |
| IGHV3-15 | *01 | *01 | *01 | *01 |
| IGHV3-16 | *02 | *02 | *02 | *02 |
| IGHV1-18 | *01 | *01 | *01 | *01 |
| IGHV3-20 | *01 | *04[a] | *01 | *04[a] |
| IGHV3-21 | *01 | *03 | *01 | *03 |
| IGHV3-23 | *01 | *04 | *01 | *04 |
| IGHV3-23D | Deleted | *01 | Deleted | *01 |
| IGHV1-24 | *01 | *01 | *01 | *01 |
| IGHV2-26 | *01 | *01 | *01 | *01 |
| IGHV4-28 | *01 | *07 | *01 | *07[b] |
| IGHV3-30 | *18 | *18 | *18 | *18 |

| | | | | |
|---|---|---|---|---|
| IGHV4-30-2 | *01 | *01 | *01 | *01 |
| IGHV3-30-3 | *01 | *03 | *01 | *03 |
| IGHV4-30-4 | Novel | Deleted | Novel | Deleted |
| IGHV3-30-5 | *01 | Deleted | *01 | Deleted |
| IGHV4-31 | Deleted | *03 | Deleted | *03 |
| IGHV3-33 | Deleted | *06 | Deleted | *06 |
| IGHV4-34 | *01 | *01 | *01 | *01 |
| IGHV3-35 | *01 | *01 | *01 | *01 |
| IGHV3-38 | *03 | *02 | *03 | *02 |
| IGHV4-38-2 | *02 | *01 | *02 | *01 |
| IGHV3-43D | *01 | Novel | *01 | Novel |
| IGHV3-38-3 | *01 | Novel | *01 | Novel |
| IGHV1-38-4 | *01 | *01 | *01 | *01 |
| IGHV4-39 | *01 | *01 | *01 | *01 |
| IGHV3-43 | *01 | *01 | *01 | *01 |
| IGHV1-45 | *02 | *02 | *02 | *02 |
| IGHV1-46 | *01 | *03 | *01 | *03 |
| IGHV3-48 | *01 | *01 | *01 | *01 |
| IGHV3-49 | *03 | *03 | *03 | *03 |
| IGHV5-51 | *01 | *03 | *01 | *03 |
| IGHV3-53 | *04 | *02 | *04 | *02 |
| IGHV1-58 | *02 | *02 | *02 | *02 |
| IGHV4-59 | *11[a] | *01 | *11[a] | *01 |
| IGHV4-61 | *09[a] | *02 | *09[a] | *02 |
| IGHV3-64 | *07[a] | *01 | *07[a] | *01 |
| IGHV3-66 | *02 | *03 | *02 | *03 |
| IGHV1-69 | *12 | *14 | *12 | *14 |
| IGHV2-70D | Deleted | *14 | Deleted | *14 |
| IGHV1-69-2 | Deleted | *01 | Deleted | *01 |
| IGHV1-69D | Deleted | Novel (1-69*05) | Deleted | Novel (1-69*05) |
| IGHV2-70 | Novel | *19[a] | Novel | *19[a] |
| IGHV3-72 | *01 | *01 | *01 | *01 |

| | | | | |
|---|---|---|---|---|
| IGHV3-73 | *02 | *02 | *02 | *02 |
| IGHV3-74 | *01 | *01 | *01 | *01 |
| IGHV7-81 | Novel | *01 | Novel | *01 |
| IGHV1-8 | *01 | *03 | *01 | *03 |
| IGHV3-9 | *01 | *03 | *01 | *03 |

[a]These alleles were novel alleles submitted to IMGT and were given allele identification prior to publication.

[b]These alleles were not detected in the parents due to decreased coverage but were present in the fosmids.

**Table S13. Sequence for novel alleles detected in NA19240**

| **Gene** | **Novel allele sequence** |
|---|---|
| IGHV4-30-4 | TCTCTGGCACAGTAATACACGGCCGTGTCTGCGGCAGTCACAGAGCTCAGCTTCA<br>GGGAGAACTGGTTCTTGGACGTGTCTACTGATATGGTAACTCGACTCTTGAGGGA<br>CGGGTTGTAGTAGGTGCTCCCACTGTAATAGATGTACCCAATCCACTCCAGGCCC<br>TTCCCTGGGGGCTGGCGGATCCAGCTCCAGTAGTAATCACCACTGCTGATGGAGC<br>CACCAGAGACAGTGCAGGTGAGGGACAGGGTCTGTGAAGGCTTCACCAGTCCTG<br>GGCCCGACTCCTGCAGCTGCAGCTG |
| IGHV3-43D | TATCTTTTGCACAGTAATACAAGGCGGTGTCCTCAGCTCTCAGACTGTTCATTTGC<br>AGATACAGGGAGTTTTTGCTGTTGTCTCTGGAGATGGTGAATCGACCCTTCACAG<br>AGTCTGCATAGTATGTGCTACCACCATCCCAACTAATAAGAGAGACCCACTCCAG<br>ACCCTTCCCCGGAGCTTGACGGACCCAGTGCATGGCATAATCATCAAAGGTGAAT<br>CCAGAGGCTGCACAGGAGAGTCTCAGGAACCCCCCAGGCTGTACCACGACTCCC<br>CCAGACTCCACCAGCTGCACTTC |
| IGHV3-38-3 | TCTTTCTTACAGTAATACACAGCCGTGTCCTCAGCTCTCAGGCTGTTCATTTGAAG<br>ATACAGCGTGTTCTTGGAATTGTCTCTGGAGATGGTGAATCTGCCCTTCCTGGAGT<br>CTGCGTAGTATGTGCTACCACCACTAATGGATGAGACCCACTCCAGACCCTTCCC<br>TGGAGCCTGGCGGACCCAGCTCATCTCATTGCTACTGACGGTGAATCCAGAGGCT<br>GCACAGGAGAGTCTCAGGGACCCCCCAGGCTGTACCAAGACTCCCCGAGACTCC<br>ACCAGCTGCACCTC |
| IGHV2-70 | GTATCCGTGCACAGTAATACGTGGCTGTGTCCACAGGGTCCATGTTGGTCATTGT<br>AAGGACCACCTGGTTTTTGGAGGTGTCCTTGGAGATGGTGAGCCTGGTCTTCAGA<br>GATGTGCTGTAGTATTTATCATCATCCCAATCAATGAGTGCAAGCCACTCCAGGG<br>CCTTCCCTGGGGGCTGACGGACCCAGCTCACACACATTCCACTAGTGCTGAGTGA<br>GAACCCAGAGAAGGTGCAGGTCAGTGTGAGGGTCTGTGTGGGTTTCACCAGCGC<br>AGGACCAGACTCCCTCAAGGTGACCTG |
| IGHV7-81 | TATCTCGCACAGTAATACATGGCCATGTCCTCAGCCTTTAGGCTGCTGATCTGCAT<br>GTATGCTATGCTGGCAGAGGTGTCCATGGAGAAGACAAACCGTCCTGTGAAGCCC<br>TGGGCATATGTTGGGTTCCCAGTGTAGGTGTTGAACCATCCCATCCACTCAAGCC<br>CTTGTCCAGGGGCCTGTGGCACCCAATTCATACCATAGGTGGTGAAACTGTAACC |

| | |
|---|---|
| | AGAAGCCTTGCAGGAGACCTTCACTGAGGCCCCAGGCTGCTTCACCTCATGGCCA GACTGCACCAGCTGCACCTG |
| IGHV1-69D | IGHV1-69*05 |

**Table S14. Alleles for IGHV genes for NA12878 and the inherited alleles in the parents of NA12878 (NA12892 and NA12891)**

| Gene | Haplotype 1 | Haplotype 2 | NA12892 | NA12891 |
|---|---|---|---|---|
| IGHV6-1 | *01 | Allele lost by VDJ | *01 | Allele not needed |
| IGHV1-2 | *04 | Allele lost by VDJ | *04 | Allele not needed |
| IGHV1-3 | *01 | Allele lost by VDJ | *01 | Allele not needed |
| IGHV4-4 | *02 | Allele lost by VDJ | *02 | Allele not needed |
| IGHV7-4-1 | *01 | Allele lost by VDJ | *01 | Allele not needed |
| IGHV2-5 | *02 | Allele lost by VDJ | *02 | Allele not needed |
| IGHV3-7 | *01 | Allele lost by VDJ | *01 | Allele not needed |
| IGHV3-11 | *01 | Allele lost by VDJ | *01 | Allele not needed |
| IGHV3-13 | *01 | Allele lost by VDJ | *01 | Allele not needed |
| IGHV3-15 | *07 | Allele lost by VDJ | *07 | Allele not needed |
| IGHV3-16 | *02 | Allele lost by VDJ | *02 | Allele not needed |
| IGHV1-18 | *01 | Allele lost by VDJ | *01 | Allele not needed |
| IGHV3-20 | *04 | Allele lost by VDJ | *04 | Allele not needed |
| IGHV3-21 | *01 | Allele lost by VDJ | *01 | Allele not needed |
| IGHV3-23 | *01 | *04 | *01 | *04 |

| | | | | |
|---|---|---|---|---|
| IGHV1-24 | *01 | *01 | *01 | *01 |
| IGHV2-26 | Novel | *01 | Novel | *01 |
| IGHV4-28 | *07 | *05 | *07 | *05 |
| IGHV3-30-3 | Deleted | *03 | Deleted | *03 |
| IGHV3-33 | Novel | Deleted | Novel | Deleted |
| IGHV4-34 | *01 | *01 | *01 | *01 |
| IGHV3-35 | *01 | Novel | *01 | Novel |
| IGHV3-38 | *02 | *02 | *02 | *02 |
| IGHV4-39 | *01 | *01 | *01 | *01 |
| IGHV3-43 | *01 | *01 | *01 | *01 |
| IGHV1-45 | *02 | *02 | *02 | *02 |
| IGHV1-46 | *04[a] | *01 | *04[a] | *01 |
| IGHV3-48 | *01 | *02 | *01 | *02 |
| IGHV3-49 | *03 | *05 | *03 | *05 |
| IGHV5-51 | *01 | *01 | *01 | *01 |
| IGHV3-53 | *02 | *01 | *02 | *01 |
| IGHV1-58 | *01 | *01 | *01 | *01 |
| IGHV4-59 | *01 | *01 | *01 | *01 |
| IGHV4-61 | *01 | *01 | *01 | *01 |
| IGHV3-64 | *02 | *02 | *02 | *02 |
| IGHV3-66 | *01 | *03 | *01 | *03 |
| IGHV1-69 | *04 | *01 | *04 | *01 |
| IGHV2-70 | *15 | *01 | *15 | *01 |
| IGHV3-72 | *01 | *01 | *01 | *01 |
| IGHV3-73 | *01 | *02 | *01 | *02 |
| IGHV3-74 | *01 | *01 | *01 | *01 |
| IGHV7-81 | *01 | *01 | *01 | *01 |
| IGHV1-8 | *01 | Deleted | *01 | Deleted |
| IGHV3-9 | *01 | Deleted | *01 | Deleted |

[a]These alleles were novel alleles submitted to IMGT and were given allele identification prior to publication.

**Table S15. Sequence for novel alleles detected in NA12878**

| Gene | Novel allele sequence |
|---|---|

| | |
|---|---|
| IGHV2-26 | GTATCCATGCACAGTAATATGTGGCTGTGTCCACAGGGTCCATATTGGTCATGGTAAGGACCACCTGGCTTTTGGAGGTGTCCTTGGAGATGGTGAGCCTGCTCTTCAGAGATGTGCTGTAGGATTTTTCGTCATTCGAAAAAATGTGTGCAAGCCACTCCAGGGCCTTCCCTGGGGGCTGACGGATCCAGCTCACACCCATTCTAGCATTGCTGAGTGAGAACCCAGAGACGGTGCAGGTCAGCGTGAGGGTCTCTGTGGGTTTCACCAGCACAGGACCAGACTCCTTCAAGGTGACCTG |
| IGHV3-33 | TCTCTCGCACAGTAATACACAGCCGTGTCCTCGGCTCTCAGGCTGTTCATTTGCAGATACAGCGTGTTCTTGGAATTGTCTCTGGAGATGGTGAATCGGCCCTTCACGGAGTCTGCATAGTATTTATTACTTCCATCATACCATATAACTGCCACCCACTCCAGCCCCTTGCCTGGAGCCTGGCGGACCCAGTGCATGCCATAGCTACTGAAGGTGAATCCAGAGGCTGCACAGGAGAGTCTCAGGGACCTCCCAGGCTGGACCACGCCTCCCCCAGACTCCACCAGCTGCACCTG |
| IGHV3-35 | TTTCTCACACAGTAATACACAGCCGTGTCCTCGGCCCTCAGGCTATTCGTTTGCAGATACAGGGTGTTCCTGGAATTGTCTCTGGAGATGATGAATTGGCCCTTCACAGAGTCTGCATAGTGCGTCCTACTGCCATTCCAACTAACACCCGATACCCACTCCAGCCCCTTTCCTGGAGCCTGATGGACCCAGTTCATGTCACTGTTACTGAAGGTGAATCCAGAGGCTGCACAGGAGAGTCTCAGGGATCCCCCAGGCTGTACCAAGCCTCCCCCAGACTCCACCAGCTGCACCTC |

**Table S16. Validation of genotyped structural variants  with fosmids and parental assemblies.**

| SV ID | Haplotype | Percent resolved* | IGHV genes | Fosmid validation | Parent validation |
|---|---|---|---|---|---|
| SV_1 | 1 | . | . | . | . |
| | 2 | 100.00% | V7-4-1 | 100.00% | 100.00% |
| SV_2B | 1 | 100.00% | V1-8, V3-9 | 99.96% | 100.00% |
| | 2 | 98.08% | V1-8, V3-9 | 100.00% | 100.00% |
| SV_3 | 1 | 100.00% | V3-23 | 100.00% | 99.95% |
| | 2 | 100.00% | V3-23, V3-23D | 100.00% | 100.00% |
| SV_4 | 1 | 100.00% | V4-28, V3-30, V4-30-2, V3-30-3, V4-30-4, V3-30-5 | 99.99% | 99.97% |
| | 2 | 100.00% | V4-28, V3-30, V4-30-2, V3-30-3, V4-31, V3-33 | 99.97% | 99.96% |
| SV_5 | 1 | 95.78% | V4-38-2, V3-43D, V3-38-3, V1-38-4 | 100.00% | 99.98% |
| | 2 | 94.41% | V4-38-2, V3-43D, V3-38-3, V1-38-4 | 100.00% | 100.00% |
| SV_6 | 1 | 96.95% | V1-69, V2-70 | 99.99% | 99.99% |
| | 2 | 100.00% | V1-69, V2-70D, V1-69-2, V1-69D, V2-70 | 99.92% | 99.96% |

*This refers to the number of bases resolved in the IGenotyper assembly relative to the custom IGH-reference.

**Table S17. Number of SNVs lifted over to GRCh37/hg19 in NA19240 and NA12878**

| Sample | IGenotyper SNVs in hg19 | IGenotyper not lifted to hg19 | 1KG Phase 3 WGS SNVs count | 1KGS Phase 3 Chip SNV count |
|---|---|---|---|---|
| NA19240 | 4474 | 703 | 3120 | 69 |
| NA12878 | 2868 | 737 | 2266 | 55 |

**Table S18. Number of overlapping SNVs in NA19240 and NA12878 between IGenotyper and the 1KGP phase 3 datasets**

| | | Count | | | Percentage | | | |
|---|---|---|---|---|---|---|---|---|
| Sample | Dataset | Overlap | IGenotyper only | Not in IGenotyper | Overlap (fraction of dataset) | Overlap (fraction of IGenotyper) | IGenotyper only (fraction of IGenotyper) | Not in IGenotyper (fraction of dataset) |
| NA19240 | 1KG WGS | 2,578 | 1,896 | 542 | 82.63% | 57.62% | 42.38% | 17.37% |
| NA19240 | 1KG Chip | 60 | 4,414 | 9 | 86.96% | 1.34% | 98.66% | 13.04% |
| NA12878 | 1KG WGS | 2,190 | 678 | 76 | 96.65% | 76.36% | 23.64% | 3.35% |
| NA12878 | 1KG Chip | 52 | 2816 | 3 | 94.55% | 1.81% | 98.19% | 5.45% |

**Table S19. Number of SNVs within accessible regions defined by 1KGP**

| Sample | Category | Total SNVs | SNVs in pilot regions | SNVs in strict regions | SNVs in in-accessible regions with pilot criteria | SNVs in in-accessible regions with strict criteria |
|---|---|---|---|---|---|---|
| NA12878 | IGenotyper only | 678 | 342 | 20 | 49.56% | 97.05% |
| NA12878 | Not in IGenotyper | 76 | 62 | 23 | 18.42% | 69.74% |
| NA19240 | IGenotyper only | 1,896 | 1289 | 165 | 32.01% | 91.30% |
| NA19240 | Not in IGenotyper | 542 | 487 | 210 | 10.15% | 61.25% |

**Table S20. Number of SNVs passing or failing ($p < 0.001$) Hardy-Weinberg equilibrium**

| Sample | Category | Total | Fail ( thres=0.001) | % Fail | % Pass |
|---|---|---|---|---|---|
| NA12878 | chr14 (no IGHV locus) | 2,607,046 | 5,793 | 0.22% | 99.78% |
| | IGHV locus | 34,393 | 2,518 | 7.32% | 92.68% |
| | Overlap with IGenotyper | 2,162 | 755 | 34.92% | 65.08% |
| | IGenotyper only | 127 | 23 | 18.11% | 81.89% |
| | Not in IGenotyper | 74 | 34 | 45.95% | 54.05% |
| NA19240 | chr14 (no IGHV locus) | 2,607,046 | 7,689 | 0.29% | 99.71% |
| | IGHV locus | 34,393 | 4,740 | 13.78% | 86.22% |
| | Overlap with IGenotyper | 2,163 | 869 | 40.18% | 59.82% |
| | IGenotyper only | 925 | 410 | 44.32% | 55.68% |
| | Not in IGenotyper | 229 | 141 | 61.57% | 38.43% |

**Table S21. Genotypes for NA19240 embedded structural variants in custom IGH reference.**

| SV ID | Genotype | IGHV gene genotypes | BioNano Support |
|---|---|---|---|
| SV_1 | ./0 | ./V7-4-1 | Yes |
| SV_2B | 0/0 | V3-9,V1-8/V3-9,V1-8 | No |
| SV_2A | 1/1 | ./. | No |
| SV_3 | 0/1 | V3-23D,V3-23/V3-23 | Yes |
| SV_5 | 0/0 | V4-38-2,V3-43D,V3-38-3,V1-38-4/V4-38-2,V3-43D,V3-38-3, V1-38-4 | Yes |
| SV_6 | 0/1 | V1-69,V2-70/V1-69,V2-70D,V1-69-2,V1-69D,V2-70 | Yes |

**Table S22. Statistics from multiplexing replicates of NA12878**

| Replicate | CCS coverage | Subread coverage | Sequence concordance | Amount of assembly recapitulated | Overlappin g variants | Missed variants | Additiona l variants |
|---|---|---|---|---|---|---|---|
| NA12878_1 | 72.98 | 660.04 | 99.97% | 98.46% | 1,706 | 429 | 45 |
| NA12878_2 | 74.11 | 666.82 | 99.99% | 98.57% | 2,029 | 56 | 48 |

| | | | | | | | |
|---|---|---|---|---|---|---|---|
| NA12878_3 | 68.64 | 623.72 | 99.99% | 98.74% | 2,027 | 58 | 49 |
| NA12878_4 | 65.58 | 594.86 | 99.99% | 98.54% | 2,061 | 24 | 41 |
| NA12878_5 | 95.37 | 819.49 | 99.99% | 93.82% | 2,052 | 34 | 41 |
| NA12878_6 | 41.30 | 358.56 | 99.99% | 95.74% | 1,972 | 112 | 15 |
| NA12878_7 | 101.62 | 872.87 | 99.98% | 94.13% | 2,042 | 44 | 66 |
| NA12878_8 | 97.73 | 837.55 | 99.98% | 94.37% | 2,048 | 38 | 49 |
| Average | 77.76 | 681.98 | 99.99% | 96.27% | 2033 | 52.29 | 44.14 |
